# The structure of HiSiaQM defines the architecture of tripartite ATP-independent periplasmic (TRAP) transporters

**DOI:** 10.1101/2021.12.03.471092

**Authors:** Martin F. Peter, Peer Depping, Niels Schneberger, Emmanuele Severi, Karl Gatterdam, Sarah Tindall, Alexandre Durand, Veronika Heinz, Paul-Albert Koenig, Matthias Geyer, Christine Ziegler, Gavin H. Thomas, Gregor Hagelueken

## Abstract

Tripartite ATP-independent periplasmic (TRAP) transporters are widespread in bacteria and archaea and provide important uptake routes for many metabolites ^1–3^. They consist of three structural domains, a soluble substrate-binding protein (P-domain), and two transmembrane domains (Q- and M-domains) that form a functional unit ^4^. While the structures of the P-domains are well-known, an experimental structure of any QM-domain has been elusive. HiSiaPQM is a TRAP transporter for the monocarboxylate sialic acid, which plays a key role in the virulence of pathogenic bacteria ^5^. Here, we present the first cryo-electron microscopy structure of the membrane domains of HiSiaPQM reconstituted in lipid nanodiscs. The reconstruction reveals that TRAP transporters consist of 15 transmembrane helices and are structurally related to elevator-type transporters, such as GltPh and VcINDY ^6, 7^. Whereas the latter proteins function as multimers, the idiosyncratic Q-domain of TRAP transporters enables the formation of a monomeric elevator architecture. Structural and mutational analyses together with an AlphaFold ^8^ model of the tripartite (PQM) complex reveal the structural and conformational coupling of the substrate-binding protein to the transporter domains. Furthermore, we characterize high-affinity VHHs that bind to the periplasmic side of HiSiaQM and inhibit sialic acid uptake *in vivo*. Thereby, they also confirm the orientation of the protein in the membrane. Our study provides the first structure of any binding-protein dependent secondary transporter and provides starting points for the development of specific inhibitors.

## Main

Tripartite ATP-independent periplasmic transporters (TRAPs) represent a structural- and functional mix of the well-studied ATP-binding cassette (ABC) transporters and secondary active transporters, by functioning as substrate binding protein (SBP) dependent secondary transporters ^1, 9–11^. They are widespread in bacteria and archaea, especially in marine environments, but absent in eukaryotic organisms. TRAP transporters, together with ABC importers and tripartite tricarboxylate transporters (TTT) ^3, 12^, define the three classes of SBP-dependent transporters. In addition to a high-affinity SBP (also named P-domain for TRAPs) that freely roams the periplasm, TRAPs consist of a smaller-(Q) and a larger-(M) membrane domain. The latter two domains are either fused into a single polypeptide chain or expressed as separate proteins that form a tight complex ^4, 13^. By far, the best-studied TRAPs are HiSiaPQM and VcSiaPQM, two sialic acid (*N*-acetylneuraminic acid) transporters from the important bacterial pathogens *Haemophilus influenzae* and *Vibrio cholerae* that are the causative agents of meningitis and cholera, respectively ^14–16^. For both pathogens, sialic acid uptake by TRAP transporters is essential for virulence and host colonization ^5, 15, 17–19^ and could provide novel targets for development of antimicrobials, for which the World Health Organization (WHO) has identified ampicillin resistant *H. influenzae* as a priority pathogen ^20^.

Of the three classes of SBP-dependent transporters, high-resolution structures for all domains are only available for ABC importers ^21^. TRAPs and TTTs have so far proved recalcitrant to experimental attempts at elucidating their structures. To date, only structures of the soluble P-domains, mainly from the family of DctP-like SBPs ^22–28^ have been determined. Features such as the role of a conserved arginine residue for high-affinity substrate binding and specificity, and the conformational rearrangement of the protein upon substrate binding have been characterized in detail ^29–31^.

Here, we determined the structure of the 70 kDa membrane domains of the HiSiaPQM TRAP transporter in lipid nanodiscs with cryo-EM. An essential step for the successful 3D reconstruction was the generation of a HiSiaQM-specific single variable domain on heavy chain (VHH) antibody and ultimately a “megabody” ^32^ that allowed us to identify HiSiaQM-filled nanodiscs during 2D classifications and aided in particle alignment. With the help of an AlphaFold ^8^ model, the structure of HiSiaQM could be confidently traced even at moderate resolution. Structural analysis of the TRAP transporter revealed high similarity to VcINDY, a dimeric elevator-type dicarboxylate transporter from *V. cholerae* ^7, 33, 34^. *In silico* modelling, mutagenesis and *in vivo* data support a new model for TRAP transporter-mediated transport, where the conformation of the SBP is coupled to the elevator movement of the transmembrane domains. Using seven VHHs that specifically bind the QM-domains, we could inhibit the TRAP transporter *in vivo*, an important step towards developing compounds that block this essential class of transporters.

## Cryo-EM structure of a TRAP transporter

We expressed the 70 kDa HiSiaQM transporter in *Escherichia coli* MC1601 cells and purified the DDM solubilized protein to homogeneity (Figure S1). For cryo-EM experiments, we reconstituted HiSiaQM in MSP1-bound DMPC lipid-nanodiscs ^35^ where helix 5 was deleted to create nanodiscs of smaller diameter with 75% of the size of native MSP1 nanodiscs ^36^. We further created a HiSiaQM-specific megabody ^32^ (Mb3) to “mark” the location of the transporter inside the nanodiscs (Figure S2). The 3D reconstruction was performed with cryoSPARC ^37^ and the resulting map had clear density for the polypeptide chain, but it was not of sufficient quality to identify side chains (Figure 1a). The advent of AlphaFold ^8^ has solved the problem of tracing the peptide chain in such low-resolution maps ^38^. We used the algorithm to create a model of HiSiaQM, which had high confidence scores (Figure S3) and could be readily placed into the reconstructed volume (Figure 1a). Slight differences between the HiSiaQM AlphaFold model and our map were adjusted with *phenix.real_space_refine* ^39^, resulting in a model-map FSC (0.143) of 6.2 Å with local resolution extending to ∼5 Å for parts of the transporter (Figure S4). The correct tracing of the chain was verified experimentally (see below). The extra density on one side of the transporter clearly corresponded to the bound Mb3 (Figure S4), unequivocally identifying the periplasmic side of the transporter (see below). The density of the megabody was not of sufficient quality to model the CDR loops with confidence, and hence it was not included in the final model (Figure 1a). We also compared our map to a previously published model of the related YiaMN TRAP transporter from Ovchinnikov et al. (∼33 % id. AA to HiSiaQM) ^40^. The model did not fit as well as the AlphaFold model, but the fold was clearly predicted correctly at the time (Figure S5 and S6).

**Fig. 1.**
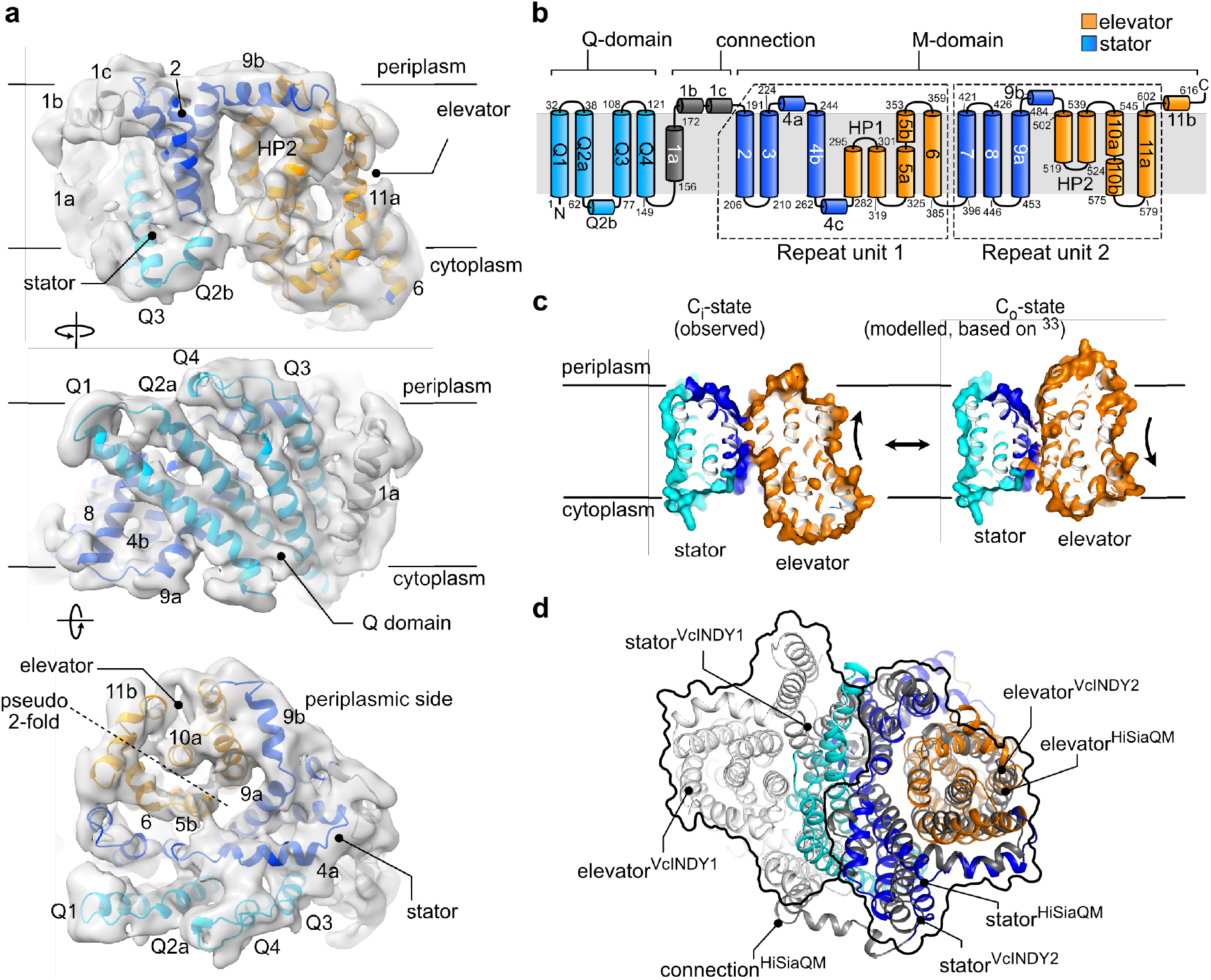
Cryo-EM structure of the TRAP transporter HiSiaQM. **a,** Structure of HiSiaQM shown as a cartoon model. The 3D reconstruction is shown in grey (contour level 0.0581). Selected secondary structure elements are indicated. The individual domains and their color code are identified in panel b. For better visualization, the density corresponding to Mb3 and the lipid belt was removed by masking operations in ChimeraX ^59^. The unmasked volume including the lipid belt-and megabody densities is shown in Figure S4. **b,** Topology diagram of HiSiaQM. **c,** Side view of HiSiaQM in the C_i_ state as determined in this work (left) and a model of the outward open C_o_ state (right) that was constructed based on the C_o_ state of VcINDY^33, 41–43^. **d,** Superposition of HiSiaQM (colored as in a) onto one unit of the dimeric VcINDY transporter in its C_i_ state (light and dark grey, PDB-ID: 5UL9). The outline of the two VcINDY proteins is shown as a black line. A more detailed view of the superposition with different orientations is shown in Figure S7.

HiSiaQM consists of 15 TM helices (Q1 - Q4 and TM1 - 11) and two helical hairpins (HP1 and HP2), which do not cross the lipid bilayer (Figure 1b). Residues 1-149 (TM helices Q1 - Q4) form the Q-domain and these four long TM helices are inclined at a 45° angle against the membrane normal and form a unique helical sheet that wraps around the M-domain. TM1 connects the Q- and M-domains and is only present in TRAP transporters such as HiSiaQM, where the Q- and M-domains are fused. The membrane topology of the transporter is shown in Figure 1b. While the periplasmic side of the transporter has a relatively flat surface, there is a deep cavity on the cytosolic side, suggesting that the transporter was captured in its inward open conformation (C_i_-state) (Figure 1c). The M-domain has an internal 2-fold pseudosymmetry that relates TM helices 2-6 and HP1 to TM helices 7-11 and HP2.

## TRAP transporters are monomeric elevators

The closest structural known homolog of the QM-domains is the VcINDY membrane transporter, a dicarboxylate transporter from *V. cholerae*, which has been demonstrated to operate via an elevator mechanism (PDB-ID: 5UL9) ^7, 33, 34^. Superposition of VcINDY onto our HiSiaQM structure illustrates that the Q-domain of HiSiaQM structurally mimics the oligomerization domain of the second polypeptide chain of the VcINDY dimer structure (Figure 1d and S7). The four helices of the Q-domain are, however, significantly longer than their structural counterparts in VcINDY. The superposition also reveals that the tips of HP1 and HP2 likely coordinate the sodium ions in HiSiaQM, which is in accordance with previous studies showing that HiSiaPQM is a symporter and translocates one sialic acid molecule together with at least two Na^+^ ions ^13^ (Figure S7).

The internal pseudo-symmetry of the elevator fold allows modeling of the outward open (Co) state of the transporter by “repeat-swap modeling” ^33, 41–43^. This procedure had been used to build a model of the outward open state of the VcINDY transporter, which was experimentally validated ^33, 44^. Using this model as a basis, we constructed a model of the outward open state of the HiSiaQM transporter by structural alignment of the elevator and stator domains (Figure 1c). Notably, the procedure did not produce any clashes in our model and agrees with the predicted tripartite complex described below. The loops connecting elevator and stator domains were adjusted manually in Coot ^45^. The C_o_-state model is available as supplemental data.

## High-affinity VHHs reveal the membrane orientation and inhibit transport

An alpaca was immunized with DDM solubilized HiSiaQM, and nine distinct VHHs were isolated, seven of which bound with nano- to picomolar affinities to at least two different regions of the transporter (Figure 2a and Figure S8 and S10). The affinity depended on the immobilization site (E235C-biotin or K273C-biotin), which can be explained by our structure, since the biotinylated residues are on different sides of the transporter.

**Fig. 2.**
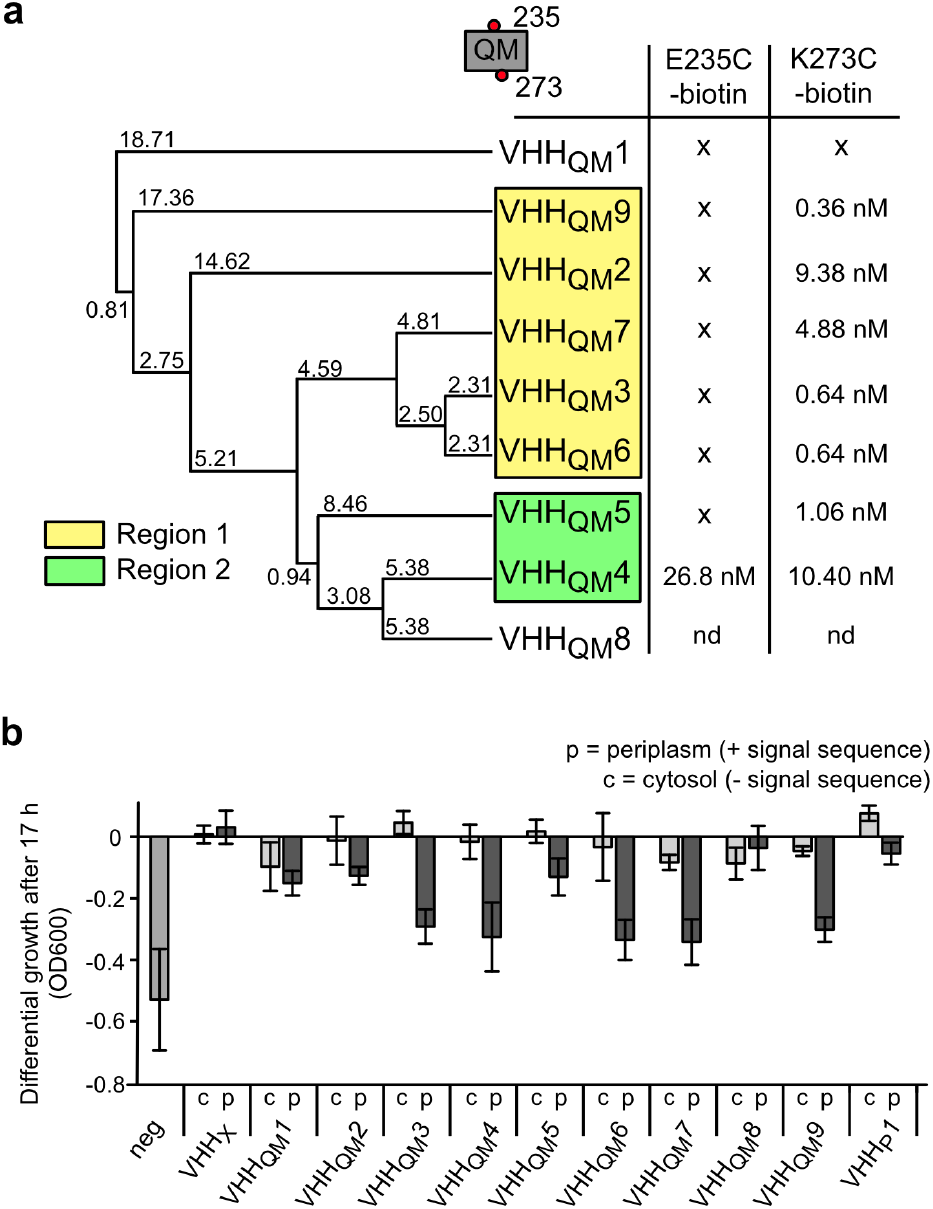
Characterization of TRAP transporter specific VHHs and inhibition of transport *in vivo*. **a,** Phylogenetic tree of nine HiSiaQM specific VHHs, calculated with Jalview ^60^. The binding of the VHHs to HiSiaQM was quantified by SPR experiments (Figure S8). Two HiSiaQM variants (E235C-biotin and K273C-biotin) were used for immobilization of the transporter on the SPR chip. VHHs that bind to HiSiaQM mutually exclusively are indicated by yellow and green boxes. The underlying data are described in detail in (Figure S10). **b,** Differential growth (i. e. growth compared to the positive control) of cultures expressing the different VHHs in the periplasm (p) or cytosol (c) in a sialic acid uptake assay. The growth of each culture was measured after 17 h.

Depending on their binding epitope, VHHs can influence the function of their target proteins, and examples of transporter-inhibiting VHHs are known, including elevator transporters ^46–49^. To test this possibility for our HiSiaQM VHHs, we used a modified *in vivo* transport assay for TRAP transporters based on *E. coli* strain SEVY3 ^13, 50^. In this strain, the native sialic acid transporter NanT was functionally replaced by HiSiaPQM, enabling it to grow in M9 minimal medium with sialic acid as the sole carbon source. We transformed different clones of the strain with our nine VHHs, either with or without a periplasmic export signal. In addition to the HiSiaQM VHHs, we included a cameloid VHH for HiSiaP (VHH_P_1) with an affinity of 0.89 µM to the SBP (Figure S9), since inhibition of SBPs by VHHs was described before ^48^. Growth curves of the different strains were recorded to investigate the effect of the VHHs on cell growth (Figure 2b). No significant inhibition of cell growth was observed when the HiSiaQM-specific VHHs were expressed without the signal sequence and thus remained in the cytosol. In contrast, when HiSiaQM-specific VHHs were exported to the periplasm, bacterial growth was inhibited by VHHs_QM_3, 4, 6, 7, and 9. Cultures with VHHs_QM_1, 8 and VHH_P_1 did show normal cell growth, irrespective of their cellular localization. This is in line with the SPR data, where no binding was detected to VHH_QM_1 and VHH_QM_8 (Figure 2a). Possible reasons for the lack of any effect of the VHH_P_1 after periplasmic export is that the VHH binds to a site of HiSiaP, which is not involved in complex formation with HiSiaQM or in substrate binding or that the high copy number of the SBP can simply not be saturated by the VHH ^51, 52^. We conclude that VHHs_QM_2, 3, 4, 5, 6, 7, and 9 bind to the periplasmic side of HiSiaQM. Since Mb3 was constructed from VHH_QM_3, this information identifies the TM orientation of our cryo-EM structure (Figure 1a).

## Constructing a model of the tripartite transport complex

Encouraged by the above-described quality of the HiSiaQM AlphaFold model, we employed the algorithm to predict the tripartite complex between HiSiaP and HiSiaQM by fusing the two proteins into a single chain. The biological rationale behind this approach was the observation that in rare cases, natural M-P fusions do occur in nature, as for example a TRAP transporter from *Acidaminococcus intestini* (Uniport ID: G4Q5D7) (Figure S11). The resulting models were very similar to each other and to the previously mentioned model by Ovchinnikov et al.^40^. In all models, HiSiaP with its closed substrate binding site (most similar to PDB-ID: 3B50^26^) is positioned in the same orientation on top of the periplasmic side of HiSiaQM (Figure 3a and Figure S11). An analysis with the PISA server ^53^ revealed that a combined surface area of ∼1980 Å^2^ is buried by complex formation. Figure 3b shows the conservation of this P-QM interface across a large number of different TRAP transporters. In the tripartite models, the N-terminal lobe of HiSiaP is bound to the stator domain of HiSiaQM, and the C-terminal lobe is bound to the elevator domain. This arrangement puts the substrate-binding cleft of HiSiaP directly on top of the presumed substrate-binding site of the transporter, as identified by superpositions with substrate bound VcINDY (Figure S7). This suggests that the N-terminal lobe of the SBP stays fixed on the stator, while its C-terminal lobe moves in concert with the up-and-down movement of the elevator during a transport cycle. Superposing the N-terminal lobe of open-state crystal structure of HiSiaP (PDB-ID: 2CEY) onto the stator domain in precisely the same manner puts its C-terminal lobe into a position that closely matches the elevator in its above-described outward open conformation (Figure 3a). The relative orientation of the P-domain to the QM-domains in our models is independently supported by superposition with TRAP SBPs that are known to form stable homodimers, for example TakP from *Rhodobacter sphaeroides* (Figure S12) ^54, 55^. The dimeric P-domains superpose such that the second monomers point upwards, not interfering with the interface of the tripartite P-QM complex.

**Fig. 3.**
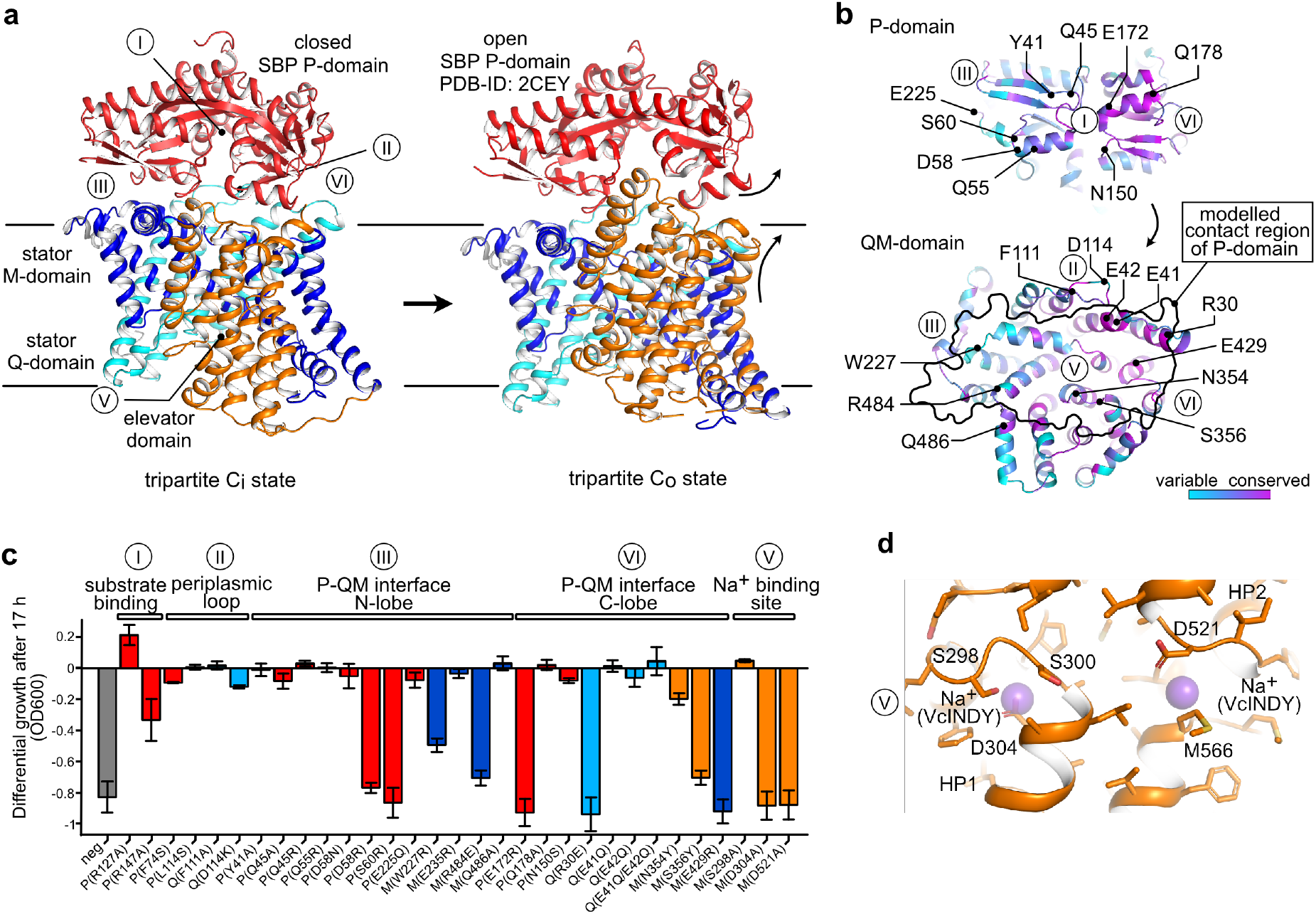
Constructing and validating the tripartite complex. **a,** Left: AlphaFold model of the tripartite complex between HiSiaQM and HiSiaP in the C_i_ state. Right: The outward open model of HiSiaQM (C_o_ state, Figure 1c). Here, the P-domain was replaced by the open (substrate-free) structure of HiSiaP (PDB-ID: 2CEY) by aligning it onto the N-terminal lobe of the P-domain in the C_i_ state (left). **b, “**Open book” view of the interface regions between QM and the N-(III) and C-lobe (IV) of the P-domain. The color code corresponds to the conservation of residues, calculated by Consurf ^61^. **c,** Differential growth (compared to a wildtype culture) of TRAP transporter mutants in a sialic acid uptake assay. The positions of the mutants with respect to the tripartite complex in panel a are indicated by labels and the color code. **d,** Presumed location of the sodium binding sites in HiSiaQM based on an alignment with the VcINDY structure shown in Figure 1d.

## Experimental validation of the tripartite model

To validate the described structures, we selected 31 residues at positions that we deemed critical for the integrity and function of the TRAP transporter, such as the substrate-binding site of the P-domain (I), an extended periplasmic loop (II), the P-QM interface (III and IV), or the assumed Na^+^ binding sites at HP1 and HP2 (V) of the QM domains (a detailed view of each mutation site is shown in Figure S13 and a sequence alignment of TRAPs in Figure S14 and S15). All mutants were tested in the above-described SEVY3-based complementation assay.

We identified ten mutants that severely or completely abolished growth and are structurally spread over all three transporter domains (Figure 3c). Mutations of highly conserved residues P^R127^ and P^R147^ in the SBP (I, Figure 3), which are known to be important for substrate binding and were previously described as a substrate-filter ^29^, showed little to no effect in the growth assay. This observation is in accordance with previous data and crystal structures of substrate-bound P^R147^ variants, where the arginine-carboxylate interaction is displaced by water molecules ^29^. It seems that at high substrate concentrations (3.2 mM in our assay) this can be tolerated by the system. Minor effects were also observed for mutants located in the long and conserved periplasmic loop connecting TMs Q3 and Q4 of the transporter or the adjacent region of the P-domain (II, Figure 3). These mutants were selected to investigate a potential “scoop-loop” mechanism, as observed in SBP-dependent ABC transporters ^56^. Twelve mutants are located in region III (Figure 3), which connects the N-terminal lobe of HiSiaP and the stator of HiSiaQM. Here, the most substantial effect on growth was observed with mutants P^D58R^ and P^S60R^. Ten of the tested mutants are located in region IV, formed by the C-terminal lobe of HiSiaP and the elevator domain of HiSiaQM. In this region, the strongest effects were seen with mutants P^E172R^, Q^R30E^, M^S356Y^, and M^E429R^. For the potential Na^+^-binding site mutants (V) two of three mutants (M^D304A^ and M^D521A^) independently led to complete loss of transporter function, providing a structural explanation for the earlier finding that a sodium ion gradient is strictly needed for transport by HiSiaPQM ^13^.

The mutants that completely inhibited growth in our uptake assay were expressed and purified to check the integrity of the protein. All QM-mutants behaved as the wildtype protein (Figure S16). For mutants located in the P-domain, the sialic acid binding properties were not affected as determined by isothermal titration calorimetry (Figure S17). Taken together, our data strongly support the validity of the tripartite model and the correct tracing of the QM-domains.

## Discussion

Our study reveals the structural architecture of TRAP transporters in a lipid bilayer. Based on our results, we suggest that TRAP transporters employ an elevator-instead of a rocker-switch mechanism. The function of the Q-domain of TRAP transporters has been elusive and our results suggest that it represents a structural mimic of the combined stator domains of multimeric elevators. It appears that a minimal size of the stator is needed to anchor the domain in the membrane and to support the up and down movement of the elevator domain. The extended length of the Q-domain helices in HiSiaQM compared to their structural counterparts in VcINDY might further stabilize this asymmetric design.

Interestingly, the translocation pathway formed by the QM-domains does not contain any conserved positively charged residues, such as Arg147 of the SBP that determines the specificity for its negatively charged monocarboxylate substrate ^29^. In contrast, the only other structurally characterized sialic acid transporter SiaT from *Proteus mirabilis* (Uniprot: HI4320) does have a conserved arginine in its translocation channel, and this residue interacts with the substrate during transport ^57^. Since SiaT does not use an SBP, it appears that the substrate selectivity of TRAP transporters is “outsourced” from the membrane domains to the SBP, explaining why the latter is strictly needed for TRAP transporter-mediated transport ^1, 4, 13^. The observation that the crystal structures of the closed- and open states of HiSiaP fit perfectly to both our inward open structure of HiSiaQM and a model of the outward open transporter, respectively, indicates that not only the interfaces of these two proteins but also their conformational flexibility have been perfectly matched by evolution.

The information presented above allows us to amend the working hypothesis for TRAP transporter-mediated transport that was last updated by Mulligan et al. in 2011 ^11^. As known from several previous studies, the SBP binds tightly (K_D_ 20-300 nM) to its substrate and switches from an opened- to a closed conformation (Figure 4, steps 1-2) ^22, 24, 31^. As the P-domain only closes when a substrate is bound, empty transport cycles are prevented ^25, 31^. In the next step, the substrate-bound SBP binds to the transporter in its inward open resting state, the structure of which has been determined in this work. Binding of the SBP must then trigger the upward movement of the elevator so that the SBP is opened and its ligand released, as has been suggested for another TRAP transporter ^58^. As the transport process is coupled to the translocation of Na^+^ ions this upwards movement is presumably accompanied by binding of two such ions (Figure 4, step 4) ^13^. Dissociation of the open SBP presumably allows the elevator domain to “fall” back into the inward open state (Figure 4, step 5). The substrate and the Na^+^ ions then diffuse into the cytosol, resetting the transport cycle (Figure 4, step 6).

**Fig. 4.**
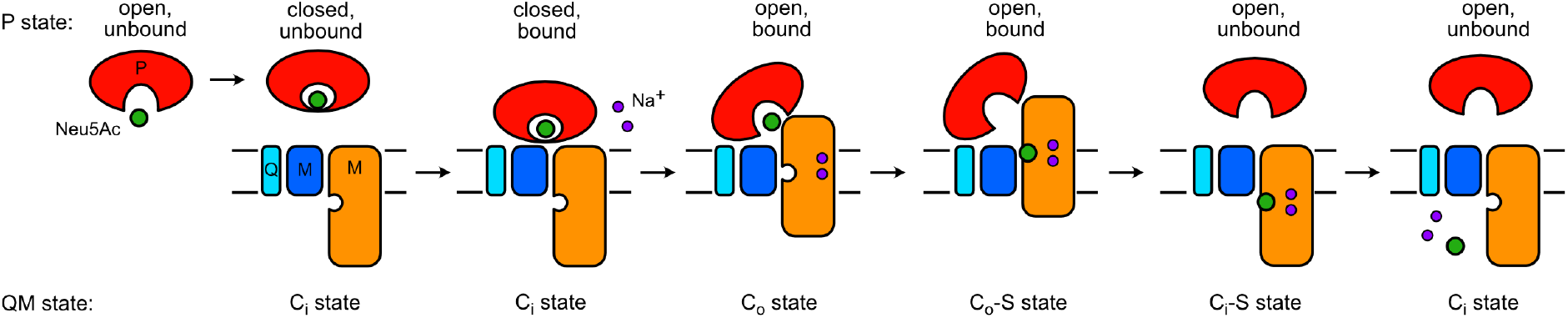
Proposed mechanism of TRAP transporters. The individual steps are explained in the main text.

Since TRAPs are absent in humans, they are attractive targets for the development of new antibiotics. Our finding that transport can be inhibited with several different VHHs opens a road into this direction. The VHHs might, for instance, disrupt the tripartite complex, or block the elevator-movement of HiSiaQM ^46^. It will be interesting to see whether small compounds can be developed that mimic the CDRs of the VHHs and thus lead to a similar effect on transport.

## Methods

### Cloning, expression and purification of HiSiaQM

As previously described in Mulligan et al. ^13^, the *HiSiaQM* gene (HI0147, Uniprot: P44543) was cloned into a pBAD vector with an N-terminal His_10_-tag and a TEV-cleavage site. For expression, the plasmid was transformed into MC1061 *E. coli* cells and a preculture of 100 mL LB-medium (100 µg/mL ampicillin) was grown overnight at 37 °C and 140 rpm. The expression was performed by inoculation of 10 L of fresh LB-medium (100 µg/mL ampicillin) with the preculture and grown for 2-3 hours at 37 °C and 140 rpm until an OD_600_ of 0.8. The protein expression was induced with 50 mg/L(+)-arabinose and incubated for 2 hours at 37 °C and 140 rpm. The cells were harvested by centrifugation at 4000 rcf for 20 min and 10 °C. The cell pellets were stored at −80 °C until further use.

For purification, the cells were resuspended in 4 times excess 50 mM KH_2_PO_4_ (pH 7.8), 200 mM NaCl and 20 % glycerol (buffer A). The cells were lysed by sonication (40 % amplitude, 5 min, pulses 10 sec on-5 sec off) on ice and afterwards centrifuged at 300,000 rcf for 1 h at 10 °C. The supernatant was discarded and the pellet was resuspended in buffer A, supplemented with 1.5 % (w/v) DDM (Dodecyl-β-D-maltoside). After incubation overnight at 4 °C under gentle shaking, the solution was again ultra-centrifuged as before. The supernatant was mixed with Ni-NTA agarose beads, which were previously equilibrated with buffer A, and incubated for 2 h under gentle shaking at 4 °C. Afterwards, the suspension was loaded onto a benchtop column at 4 °C, the flowthrough was discarded and the beads were washed with 100 mL 50 mM KH_2_PO_4_ (pH 7.8), 200 mM NaCl and 0.035 % DDM (buffer B) with 22 mM imidazole. The protein was eluted with 15 mL buffer B, supplemented with 250 mM imidazole and concentrated to 0.5 mL with a Vivaspin (100 kDa MWCO). The protein was loaded onto an equilibrated Superdex 200 increase 10/300 column on an ÄKTA chromatography system and eluted in fractions from the column with buffer B. All purification steps and the fractions from the size-exclusion chromatography were checked with an SDS-PAGE and HiSiaQM containing fractions were combined, concentrated to around 15 mg/mL and flash-frozen for storage at −80°C.

### Cloning, expression and purification of HiSiaP

The SBP was previously cloned into a pBADHisTEV vector which fuses the HiSiaP protein to an N-terminal His_6_-tag and TEV-cleavage site. The protein was expressed and purified as described in Peter et al. ^25^ for the homologous protein VcSiaP from *V. cholerae*. The protein was expressed in M9 minimal medium to avoid contamination with sialic acid and the His-tag was removed in an overnight digestion with TEV protease.

### Expression and purification of MSP1D1-H5

The *msp1d1* gene was ordered from Addgene (#20061) in a pET28a vector which fused an N-terminal His_6_-tag and TEV-cleavage site to the MSP1 protein. To decrease the size of the nanodiscs, helix 5 (residues L68-L87) was deleted via PCR ^62^, yielding construct MSP1D1-H5^36^. The expression and purification were based on ^63^. In brief, the plasmid was transformed into *E.* coli BL21 cells and a preculture of 120 mL LB-medium (50 µg/mL kanamycin) was grown for 4-6 h at 37 °C and 140 rpm until an OD_600_ of 0.6-0.8. The culture was stored at 4 °C overnight and used to inoculate 6 L TB medium (50 µg/mL kanamycin) on the next day. The culture was incubated at 37 °C and 140 rpm until an OD_600_ of 2-5-3.0 and expression was induced with a final concentration of 1 mM IPTG in the culture. After expression of 1 h at 37°C followed by 3.5 h at 28 °C under shaking with 140 rpm, the cells were harvested by centrifugation at 4000 rcf for 15 min. The cell pellet was stored at −80 °C until further use.

For purification of MSP1D1-H5, the cell pellet was resuspended in 2.5 times excess of lysis buffer (20 mM NaPi (pH 7.4), 1 % Triton X and 0.001% nuclease). Cells were lysed by sonication for 5 min at 40 % amplitude with pulses of 10 sec on, 5 sec off, under constant cooling on ice. After centrifugation of the lysate for 30 min at 30,000 rcf and 10 °C, the supernatant was mixed with Ni-NTA agarose beads, previously equilibrated with basic buffer (40 mM Tris (pH 8.0), 300 mM NaCl) and incubated for 1 h at room temperature under gentle shaking. The suspension was transferred onto a bench-top column at room temperature and the flowthrough was discarded. The beads were washed in four steps of 12 mL with basic buffer, supplemented in the first step with 1% Triton X, in the second step with 50 mM sodium-cholate and no supplementation in the third step. Afterwards, the protein was eluted from the column with 15 mL basic buffer with 300 mM imidazole and concentrated to a total volume of 5 mL. The eluate was dialyzed overnight at 4 °C against 100 times excess of buffer with 40 mM Tris (pH 8.0). For further purification, the protein solution was loaded onto a HiLoad Superdex 200 16/600 column on an ÄKTA chromatography system, previously equilibrated with 10 mM Tris (pH 7.4), 100 mM NaCl and 1 mM EDTA. The protein was eluted in fractions and checked via an SDS-PAGE for protein containing fractions. Selected fractions were combined and incubated overnight at 4 °C with TEV protease in a 1:50 ratio. Afterwards, the cleaved tag and TEV protease were removed via Ni-NTA affinity chromatography and cleaved MSP1D1-H5 was collected in the flowthrough. TEV-cleavage was checked via SDS-PAGE and the protein was concentrated and stored at −80 °C.

### Reconstitution of HiSiaQM in MSP1D1-H5 nanodiscs

The nanodisc reconstitution procedure for the TRAP transporter was based on ^63^. The purified MSP1D1-H5 and HiSiaQM were mixed in buffer C (50 mM KH_2_PO_4_ (pH 7.8), 200 mM NaCl), supplemented with 50 mM DMPC and 100 mM sodium cholate, to a ratio of 1:80:0.2 (MSP1D1-H5:DMPC:HiSiaQM). The solution was diluted with buffer C to a final concentration of 11 mM DMPC and 22 mM sodium-cholate and incubated for 2 h at 26 °C under gentle shaking. Afterwards, the reconstitution mix was dialysed in a tube with 6-8 kDa MWCO against 500 mL buffer C at 26 °C for 16 h with 4 buffer exchanges.

### Generation, cloning, expression and purification of VHHs

VHHs were generated by the Core Facility Nanobodies of the University of Bonn. One alpaca (Vicugna pacos) was immunized by six subcutaneous injections over 12 weeks with 200 µg DDM detergent-solubilized wild-type HiSiaQM mixed 1:1 (v:v) with GERBU-FAMA adjuvant. After the immunization peripheral blood mononuclear cells (PBMCs) were isolated from 100 mL blood. The RNA was extracted from PBMCs, reversely transcribed to cDNA. VHH sequences were amplified by PCR using specific primers and cloned into a phagemid vector. *E. coli* TG1 cells were transformed with the VHH phagemid library and infected with helper phage VCSM13 to produce phage displaying VHHs as pIII fusion proteins. VHHs were enriched by phage panning on HiSiaQM E235C-biotin and K273C-biotin (see below) bound to streptavidin beads. *E. coli* ER2738 were infected with the enriched phages and after one round of panning, individual clones were grown in 96-well plates. VHHs in the supernatants were tested for specific binding by ELISA. Candidates were sequenced and grouped by sequence similarity. The HiSiaP-VHH was identified during phage display with a VHH library from a homologous VcSiaP immunization campaign with the same procedure as described above.

For protein expression, the genes were cloned into a pHEN6 vector which fused the protein to an N-terminal pelB signal peptide and C-terminal LPETG sortase motif and a His_6_-tag. The plasmids were transformed into *E. coli* WK6 cells and 1 L of TB-medium (100 µg/mL ampicillin) was inoculated with 25 mL of a LB-medium (100 µg/mL ampicillin) preculture. The culture was grown at 37 °C and 140 rpm until a cell density of 0.6 and induced with a final concentration of 1 mM IPTG in the medium. The protein expression was performed overnight at 30 °C and 140 rpm and the cells were harvested by centrifugation at 5000 rpm for 20 min. The cell pellets were stored at −80 °C until further use.

For lysis of QM-VHHs, the pellet was resuspended in 15 mL extraction buffer containing 200 mM Tris (pH 8.0), 0.65 mM EDTA and 500 mM sucrose and incubated for 1 h under gentle shaking at 4 °C. The solution was treated with 70 mL of 0.25 times diluted extraction buffer and incubated overnight at 4 °C under gentle shaking for osmotic lysis. Afterwards, the solution was centrifuged at 8000 rcf for 40 min and the supernatant was filtered through a 0.45 µm filter. For the HiSiaP-VHH, the cells were lysed by sonication (40% amplitude, 5 min, pulses 10 sec on-5 sec off), centrifuged for 20 min at 20 000 rcf and 4 °C, and also filtered through a 0.45 µm filter.

For both types of VHHs, the soluble fraction after lysis was filtered and mixed with Ni-NTA beads, previously equilibrated with 0.25 times diluted extraction buffer, and incubated for 1 h at 4 °C under gentle shaking. The suspension was loaded onto a bench-top column at room temperature, the flowthrough was discarded and the column was washed with 50 mL wash buffer (50 mM Tris (pH 7.5), 150 mM NaCl, 10 mM imidazole). Subsequently, the protein was eluted from the column with 10 mL elution buffer (50 mM Tris (pH 7.5), 150 mM NaCl, 500 mM imidazole) and concentrated to 5 mL with a Vivaspin (MWCO 3 kDa). The solution was loaded onto a HiLoad Superdex 75 16/600 on an ÄKTA chromatography system, equilibrated with 10 mM Tris (pH 7.3) and 140 mM NaCl for QM-VHHs and 50 mM Tris (pH 8) and 50 mM NaCl for the HiSiaP-VHH. The eluted fractions were checked on an SDS-PAGE, protein containing fractions were combined, concentrated, flash-frozen and stored at −80 °C.

### Cloning, expression and purification of a megabody

The gene design and cloning procedure for a HopQ megabody was based on Uchański et al. ^32^. The *hopQ* gene was ordered at Addgene and cloned into the pHEN6 vector to yield the same fused tags as the VHHs. Afterwards the VHH gene of VHH_QM_3 (without the LPETG-sortase motif) was cloned into the megabody sequence via two SapI restriction sites, yielding the HopQ megabody (Mb3). The expression, lysis and purification of Mb3 were performed as described for the VHHs. For the final purification step, a HiLoad Superdex 200 16/600 column was used.

### Site-specific biotinylation of HiSiaQM

For specific biotinylation of HiSiaQM, QuickChange mutagenesis after Liu et al. ^62^ was used to design a non-cysteine HiSiaQM construct (C94A, C325S, C334S, C400S and C458S) in which cysteines (E235C or K273C) were introduced. Expression and purification were performed as described for the wildtype protein above. The labelling was performed during Ni^2+^-affinity chromatography of the HiSiaQM purification process. After 100 mL wash of the column with buffer B, supplemented with 22 mM imidazole, the column was washed with 50 mL buffer B with 1 mM TCEP, and again with 50 mL buffer B. The protein was eluted with 15 mL buffer B with 250 mM imidazole and supplemented with 100 times excess of a biotin-maleimide label (Maleimide-PEG2-biotin (EZ-Link^TM^), ThermoFisher Scientific). The solution was incubated overnight at 4 °C and further purified on an Superdex increase 200 10/300 as described previously.

### SPR VHH/megabody binding characterization

The SPR experiments were performed on a Biacore^TM^ 8K instrument (GE healthcare life sciences), using a streptavidin-functionalized sensor chip (Serie S Sensor SA, GE healthcare life sciences) and HiSiaQM buffer B at 25 °C chip temperature. The two biotinylated HiSiaQM constructs, E235C-biotin and K273C-biotin, were injected and immobilized on the chip (100 nM, 5 µL/min, 70-100 s,). The analytes, the VHHs or the megabody, were added in six injections (30 µL/min, 120 s) with doubling of the concentration at each step, resulting in single-cycle kinetic titration curves. The binding data were double referenced by blank cycle and reference flow cell subtraction. For epitope binning, the VHHs were tested for competitive binding using a ABA-injection protocol. These binding curves were fitted and analysed using the Biacore Insight Evaluation Software (version 2.0.15.12933).

### ITC binding experiments

ITC experiments were performed as described before in Peter et al. ^25^ on a MicroCal PEAQ device from Malvern Panalytical, using corresponding MicroCalPEAQ-ITC software (version 1.21) for experiment design, measurement and analysis. Sialic acid (Carbosynth) was dissolved in the P-domain standard buffer (50 mM Tris, 50 mM NaCl (pH 8)). Each mutant was at least measured three times and the mean values are given.

### Cryo-EM – Sample preparation, data collection, processing, structure modelling

The cryo-EM structural studies were performed with HiSiaQM in MSP1D1-H5 nanodiscs with DMPC lipids. For selection of HiSiaQM-containing nanodiscs, the purified Mb3 was loaded onto Ni-NTA beads, equilibrated with HiSiaQM buffer C, for 1 h at 4 °C. The flowthrough was discarded and the column was washed with 10 mL buffer C. The Mb3 bound Ni-NTA beads were resuspended in 2 mL buffer C, transferred into a flask and supplemented with HiSiaQM-nanodisc reconstitution mix after dialysis. The mixture was incubated for 1 h at 4 °C and transferred back to the bench-top column. The flowthrough was discarded, the column was washed with 10 mL buffer C and the protein was eluted with 1.5 mL buffer C with 500 mM imidazole. After concentration to around 50 µL in a Vivaspin (MWCO 100 kDa), the protein solution was loaded onto a Superose increase 6 3.2/300 column on an Agilent HPLC 1260 infinity II, previously equilibrated with buffer D (20 mM Tris (pH 7.5) and 100 mM NaCl). The eluted protein was monitored and fractionated manually and dropwise. The fractions that corresponded to the main protein peak were combined and used for cryo-EM experiments.

The grid preparation was performed on a Vitrobot (ThermoFisher Scientific) at 4 °C and 100 % humidity by using Quantifoil R1.2/1.3 grids. The blot time was set to 6 second and the blot force to 0. The grids were stored in liquid nitrogen until data collection at a Titan Krios microscope (Table S1). A dataset of 5000 movies was collected using a Krios Titan microscope equipped with a Gatan K3 camera at 300 kV. After patch motion- and CTF-correction with cryoSPARC ^37^, 2.3 million particles were automatically picked using a blob-picker job and subjected to multiple rounds of 2D classification. Representative 2D classes are shown in Figure S4 and clearly show the megabody-HiSiaQM complex in the top views. 973K particles were used to calculate 3 ab-initio structures with cryoSPARC. After heterologous-refinement and non-uniform refinement of each class, the transporter was visible in the Coulomb potential maps at a GSFSC resolution of ∼6.2 Å with a local resolution of the transporter up to ∼5 Å. To improve the map, we created a mask around the transporter/Mb3 complex and subtracted the nanodisc density outside this mask from the particles. Local non-uniform refinement with the subtracted particles yielded an improved map with GSFSC resolution of ∼4.5 Å.

### TRAP transporter *in vivo* growth assay

The growth assay was performed in modified *E. coli* cells (SEVY3 cells) without their native sialic acid nanT (ΔnanT) transporter ^50^. SEVY3 is a derivative of BW23115Δ*nanT* ^13^ carrying two further unmarked, in-frame deletions of the repressor genes, *nanR* and *nagC*, respectively controlling the expression of the NanATEK-YhcH and NagAB branches of the sialic acid utilization pathway in *E. coli* ^64–66^. This strain was obtained by sequential allelic replacement of the WT genes with “scar” sequences (coding for 19-residue-long “vestigial” peptides), made by SOEing PCR and cloned in the conditional counter-selectable vector, pKO3 ^67^, which were then cycled through the parental strain using the method outlined in the same reference. SEVY3 was confirmed by PCR and sequencing of all three deletion loci.

A plasmid (pES7^13^) containing an ampicillin resistance and the genes for HiSiaP and HiSiaQM, was transformed into competent SEVY3 cells, as described before in Mulligan et al. ^13^. If necessary, mutations were previously introduced via QuickChange mutagenesis ^62^ and checked via sequencing. For co-transformation and co-expression of VHHs in SEVY3 cells, the VHH genes were amplified with or without pelB signal sequence and cloned into a pET28a vector with kanamycin resistance. The VHH containing plasmids were transformed into competent SEVY3 cells which contained the HiSiaPQM plasmid. For VHH-supplemented growth assays, the positive control with HiSiaPQM and without VHH was co-transformed with an empty pET28a vector.

For the TRAP transporter growth assay, the SEVY3 cells with the transformed plasmids were grown in 5 mL LB-medium with corresponding antibiotics at 37 °C and 140 rpm overnight. The cells were harvested at 4000 rcf for 15 min, the supernatant was discarded and the pellet was resuspended in 10 mL M9 minimal medium. The cell suspension was centrifuged with similar conditions and the washing procedure was repeated. After a third similar centrifugation step, the pellet was resuspended in 5 mL M9 minimal medium. 1 mL of this washed culture was used to inoculate 50 mL M9 minimal medium, supplemented with 1 mM IPTG, 2 mM MgSO_4_, 0.1 mM CaCl_2_, 0.1% (w/v) sialic acid (Carbosynth) and corresponding antibiotics. The cultures were grown for 17 hours at 37 °C and 140 rpm in baffled flasks and the cell density was regularly measured on a NanoDrop 2000 (ThermoFisher Scientific) at 600 nm.

### Structural predictions with AlphaFold (deepmind)

The source code of the AlphaFold (deepmind) algorithm was downloaded from https://github.com/deepmind/alphafold and installed as described https://github.com/deepmind/alphafold. The pLDDTs scores were extracted and mapped onto the structures with PyMOL (Figure S3).

## Acknowledgements

This project was financed by the German Research Foundation (Deutsche Forschungsgemeinschaft, DFG) in projects no. HA 6805/4-1 and HA 6805/5-1 and a Method Development Grant by the TRA Life and Health (University of Bonn) as part of the Excellence Strategy of the federal and state governments (to GH and PAK). MFP acknowledges a PhD fellowship from the Konrad Adenauer Stiftung. We thank Elif Tokmak for technical assistance with cloning of the HiSiaQM mutants. We thank Florian I. Schmitz and Jan Tödtmann of the Core Facility Nanobodies (CFN) at the University of Bonn for their support. This work benefited from access to the Integrated Structural Biology platform of the Strasbourg Instruct-ERIC center IGBMC-CBI (Institut de génétique et de biologie moléculaire et cellulaire – Centre de Biologie Intégrative) Illkirch-Graffenstaden (FR). We thank Nils Marechal and Corinne Crucifix of the IGBMC-CBI for their help with sample preparation and data collection. MFP and GH thank the group of CZ (University of Regensburg) for initial cryo-EM experiments and the group of Prof. Dr. Bettina Böttcher (University of Würzburg) for preliminary cryo-EM data collections. GHT & ES thank UKRI BBSRC for funding much initial work on this system & the BBSRC White Rose DTP for a studentship to ST & Prof. Phill Stansfeld (University of Warwick, UK) for building another initial model of SiaQM. MFP and GH thank Janine Glaenzer and Prof. Dr. Olav Schiemann (University of Bonn) for support during the initial stages of this project.

## Author contributions

GH acquired funding and supervised this study. All proteins, were expressed and purified by MFP except the HiSiaP-VHH, which was provided by NS. PD and MFP developed the reconstitution of the nanodiscs. PAK supervised alpaca immunization, selection and initial characterization of VHHs. Labelled mutants for SPR experiments were prepared by MFP. KG performed and analysed the SPR experiments. Cryo-EM experiments were performed by MFP, GH and AD. The cryo-EM dataset was processed by MFP and GH. The results were discussed with CZ, VH and AD. The TRAP transporter complementation assay was elaborated and performed by MFP with the help of ES who developed the original assay and k/o strain. ITC experiments were performed by MFP and VHH_P_1 measurements by NS. MFP and GH wrote the manuscript together with GHT. All authors discussed the data and commented on the final manuscript version.

## Supporting Information

**Fig. S1.**
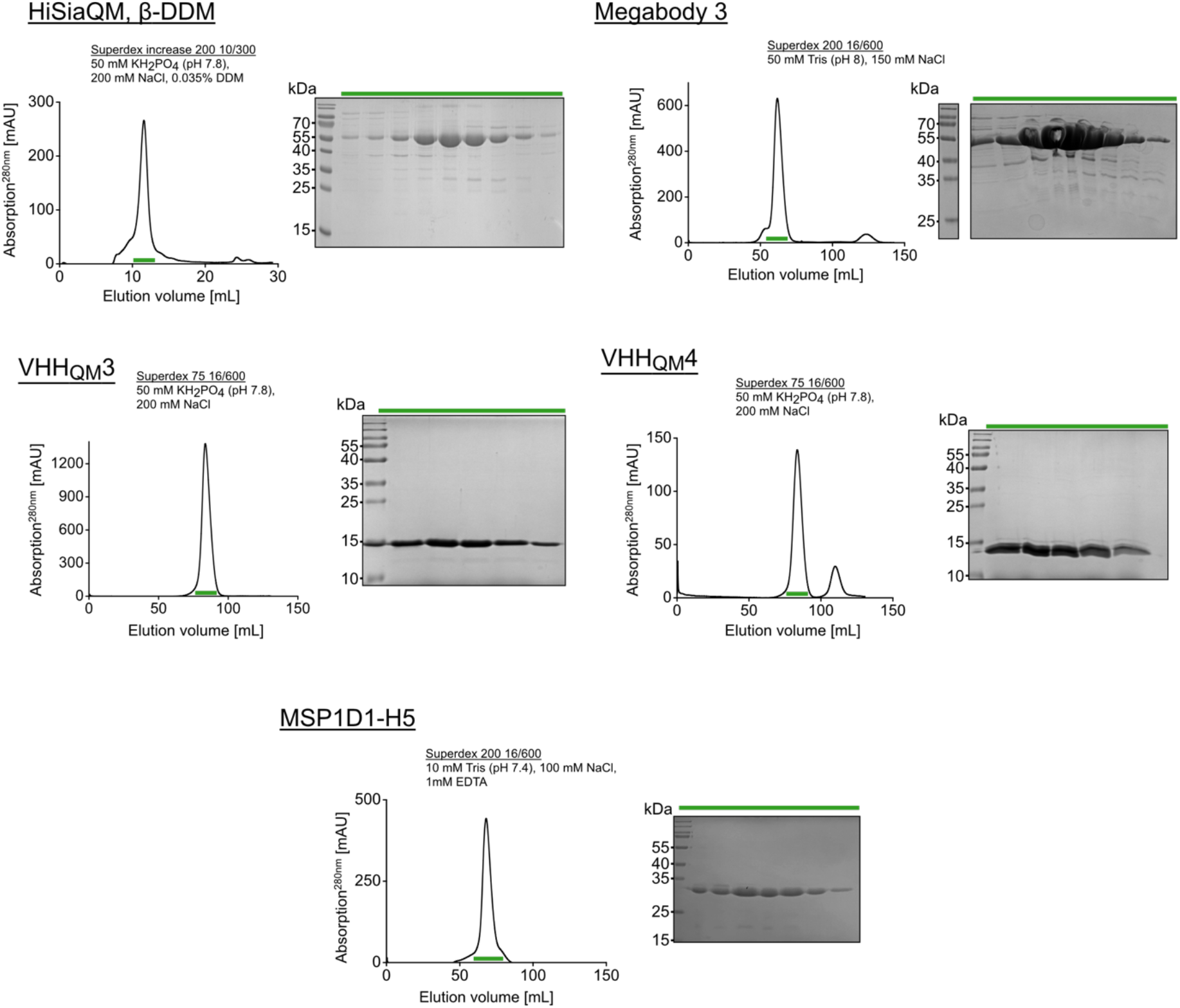
Quality of proteins used in this study. A size-exclusion chromatography profile is shown for each protein. The buffers and SEC columns are given next to the chromatograms. The fractions highlighted in green were analysed by SDS-PAGE, shown on the right side of the corresponding SEC profile.

**Fig. S2.**
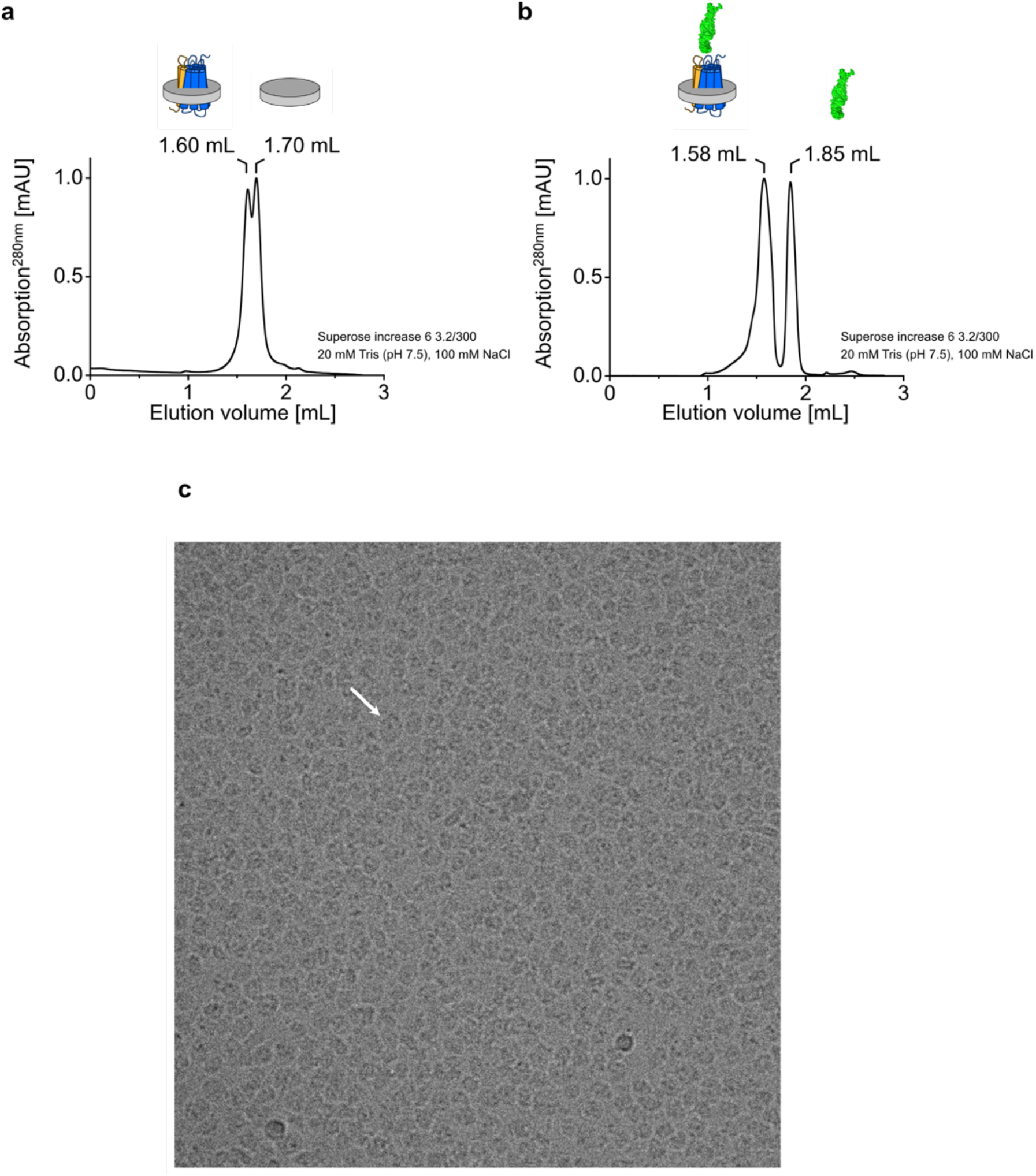
Cryo-EM sample preparation and raw micrograph of the HiSiaQM/Mb3 complex in MSP1D1-H5 nanodiscs. **a,** Normalized SEC elution profile of the HiSiaQM-MSP1D1-H5 reconstitution mixture. **b,** Normalized SEC elution profile of the HiSiaQM-MSP1D1-H5 reconstitution mixture after Ni-NTA affinity chromatography pull-down via immobilized Mb3. **c,** The picture was recorded with a ThermoFisher Glacios microscope equipped with a K2 camera at 200 kV. The white arrow marks a nanodisc in top-view with Mb3 visible as a dark spot.

**Fig. S3.**
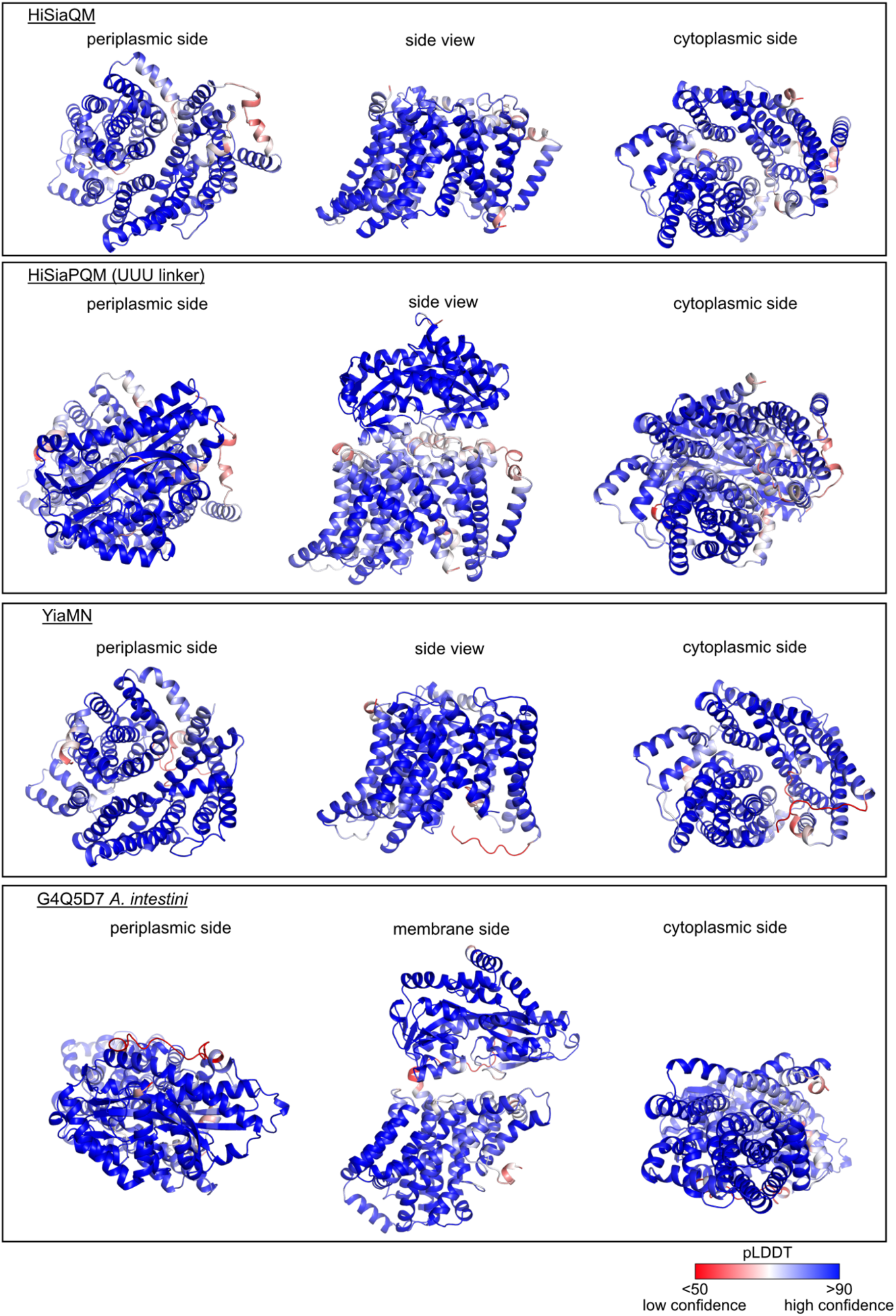
AlphaFold models that were used in this study. The pLDDT confidence score is indicated by a color gradient.

**Fig. S4.**
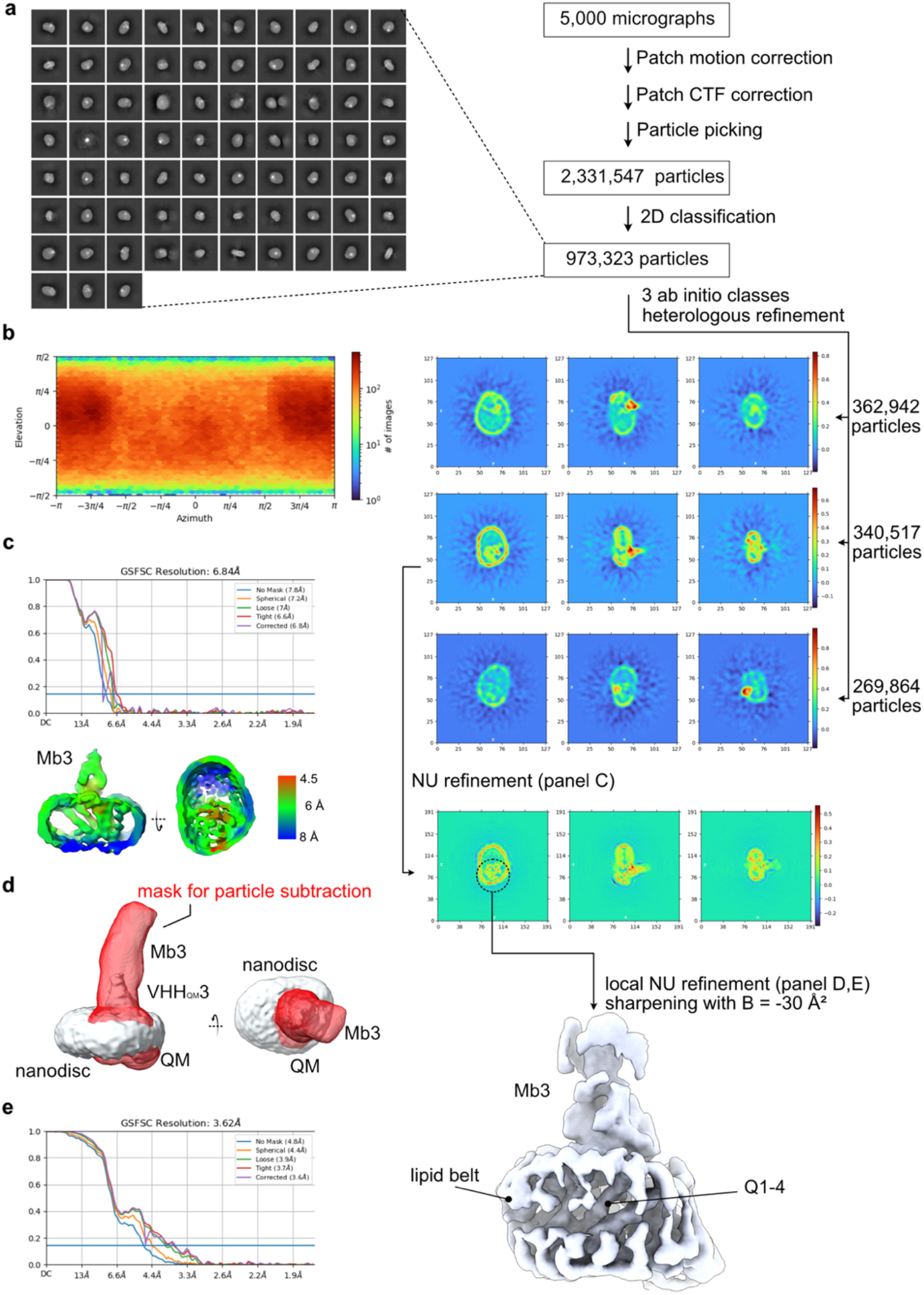
Cryo-EM processing workflow. **a,** CryoSPARC workflow for the 3D reconstruction of HiSiaQM in lipid nanodiscs with a megabody bound to the periplasmic side. **b,** Distribution of viewing angles. **c,** Gold-standard FSC curves for the NU refinement step before particle subtraction and local resolution plots (FSC =0.143). The latter are cutaways to visualize the transporter inside the nanodisc. **d,** Two views of the mask that was used for particle subtraction (red surface) relative to the refined volume. **e**, Gold-standard FSC curves for the local NU refinement after particle subtraction.

**Fig. S5.**
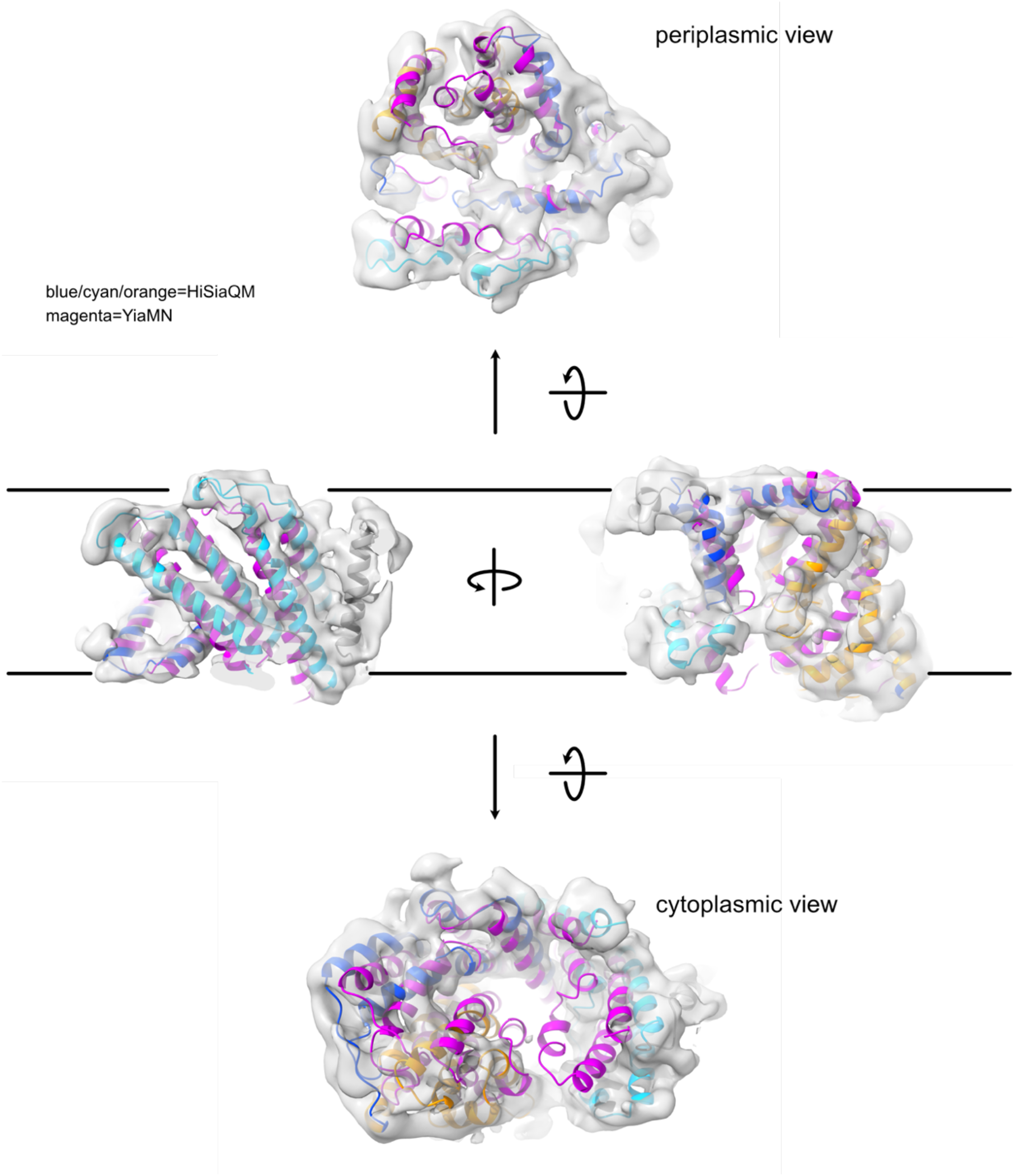
Fit of different TRAP transporter models to the cryoEM density. The YiaMN model (magenta) is from Ovchinnikov et al. ^1^. The HiSiaQM model was determined in this study with the color code as defined in Figure 1.

**Fig. S6.**
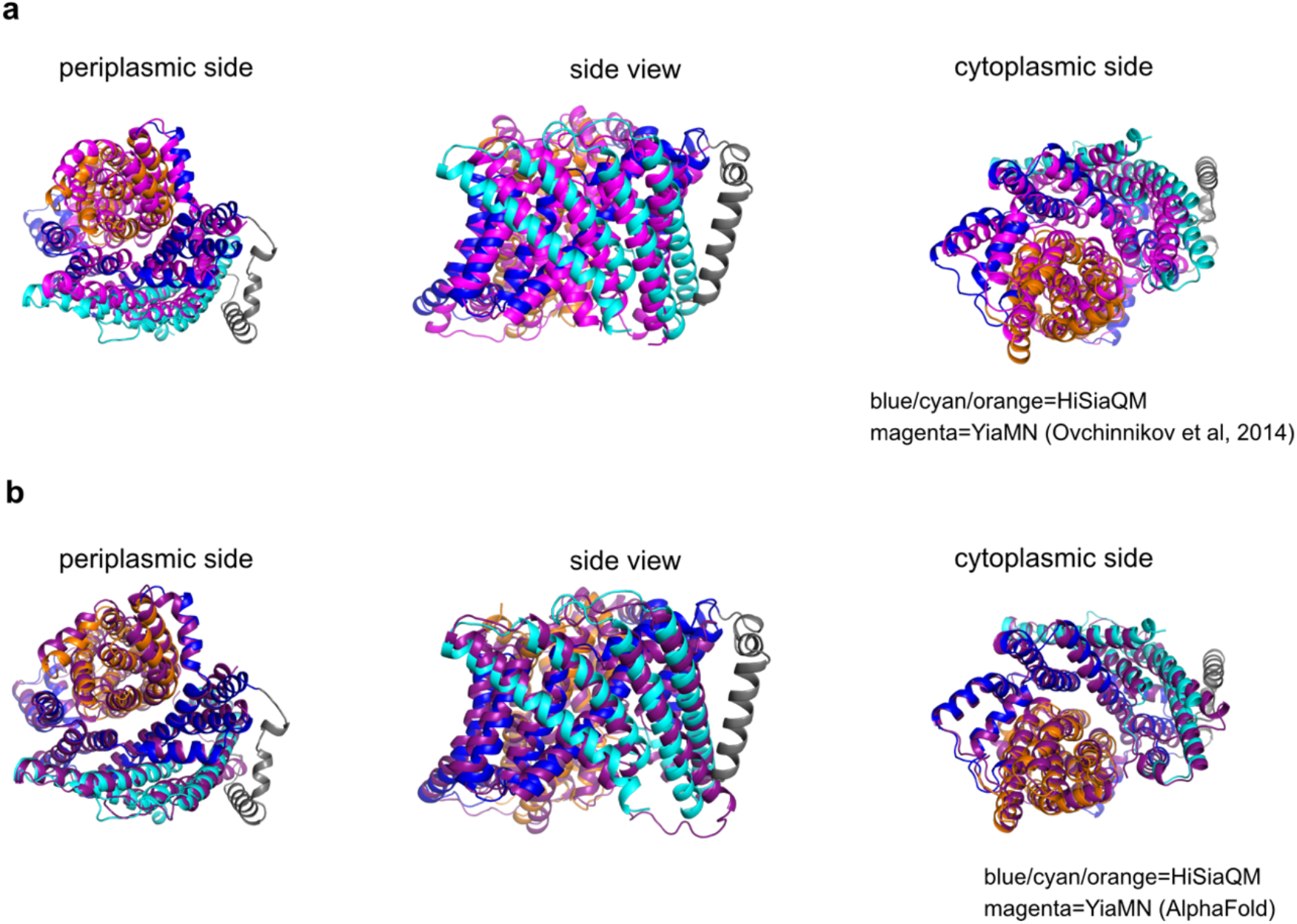
Alignment of modelled TRAP transporters. **a,** Alignment of HiSiaQM structure to the model of YiaMN from Ovchinnikov et al. ^1^. **b,** Alignment of HiSiaQM structure to the model of YiaMN from AlphaFold ^2^.

**Fig. S7.**
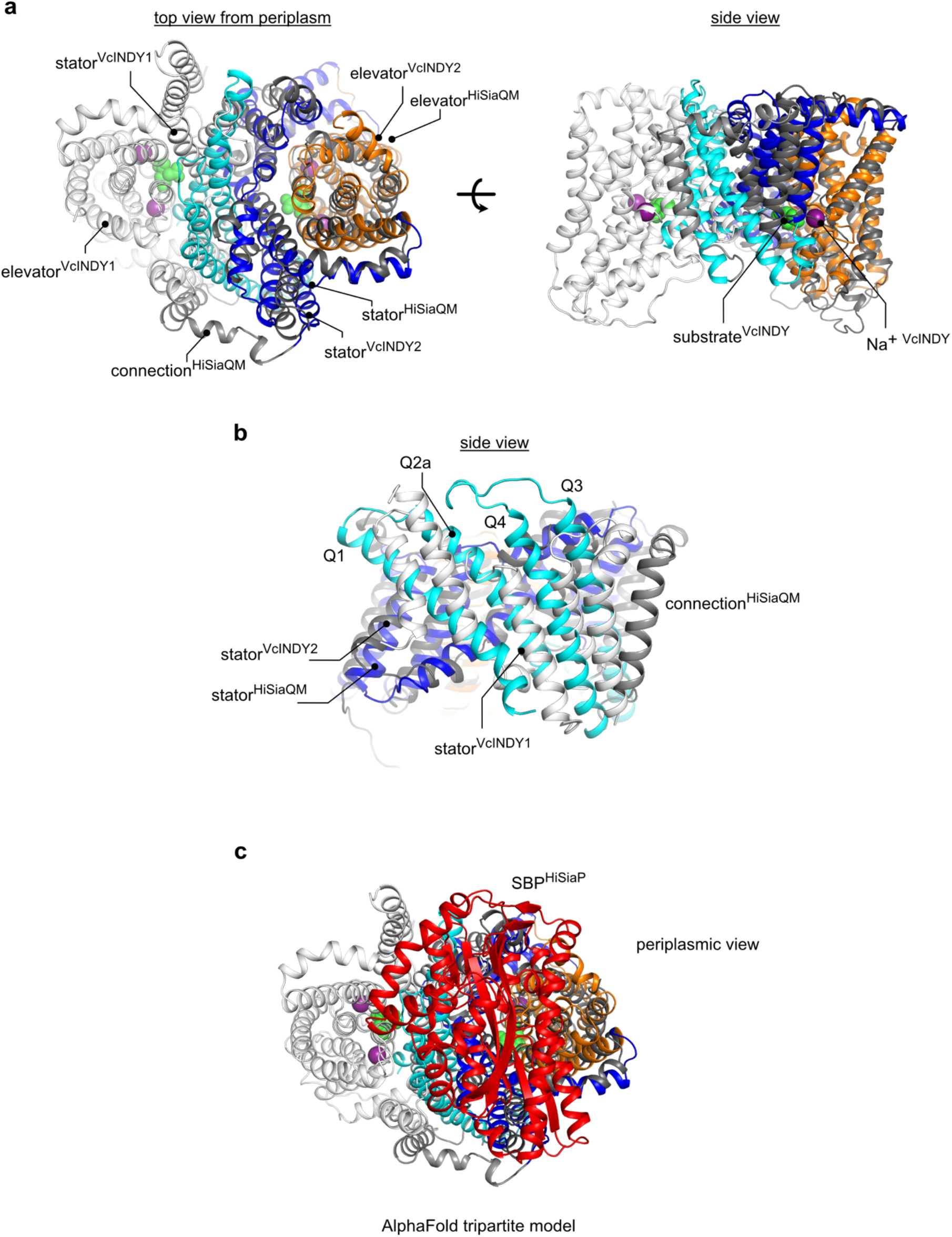
Alignment of the TRAP transporter HiSiaPQM with VcINDY. **a,** Alignment of the HiSiaQM structure (colored as in Figure 1) onto one monomeric unit of the VcINDY structure (white). The bound Na^+^ ions (purple) and citric acid (green) in each VcINDY monomer are represented as spheres. **b,** Focused view of the alignment in panel a on the Q1-Q4 helices. **c,** Same alignment as in panel a, but with the tripartite HiSiaPQM AlphaFold model (colored) from Figure 3 to VcINDY (white). The HiSiaP SBP is shown in red.

**Fig. S8.**
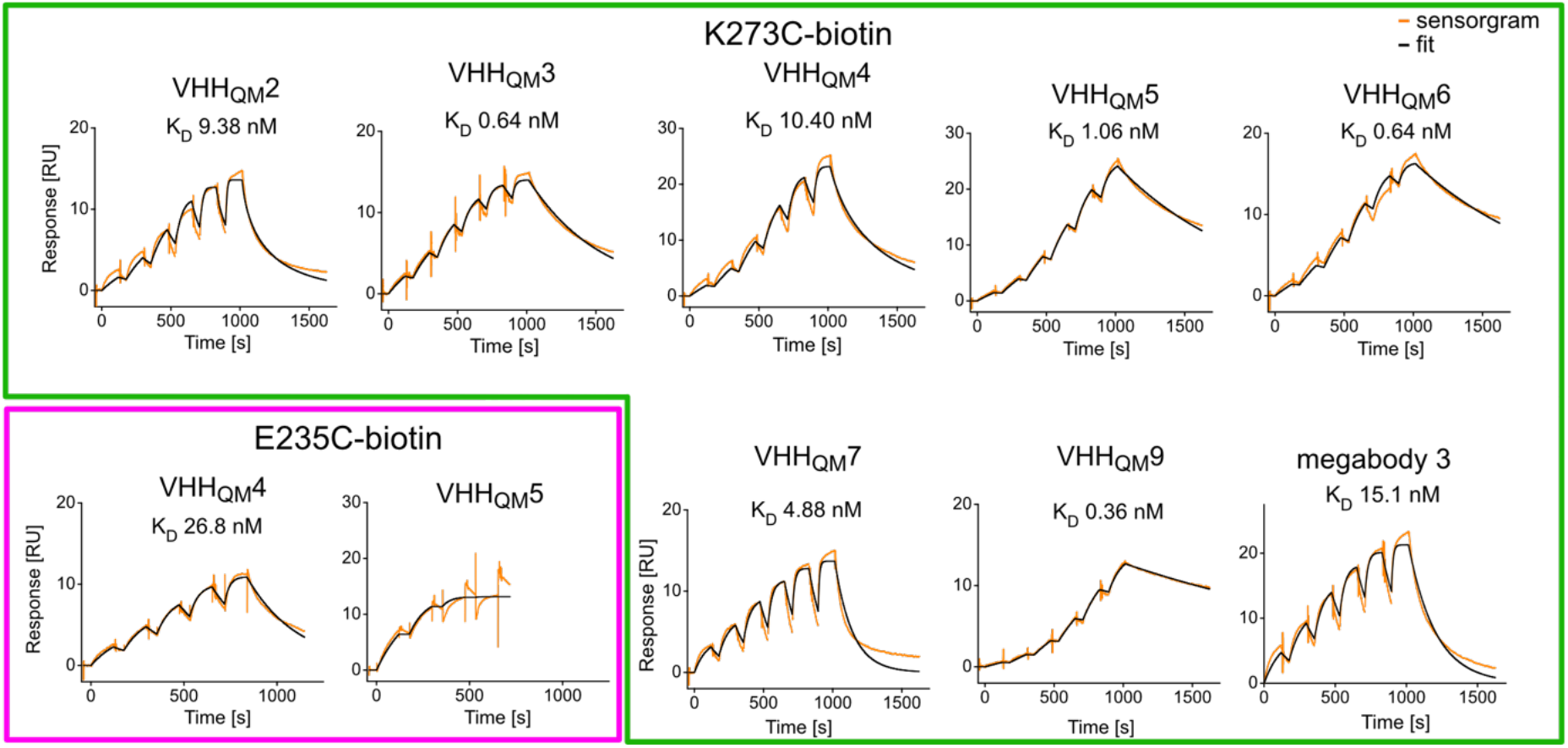
Characterization of VHH binding via SPR. Single cycle kinetic binding data for TRAP transporter VHHs. Two different immobilized HiSiaQM variants, E235C-biotin (magenta) and K273C-biotin (green), were used. The experimental data are shown in orange and the fit in black. The binding affinities are mentioned next to the sensorgram, the kinetic parameters are summarized in Table S2.

**Fig. S9.**
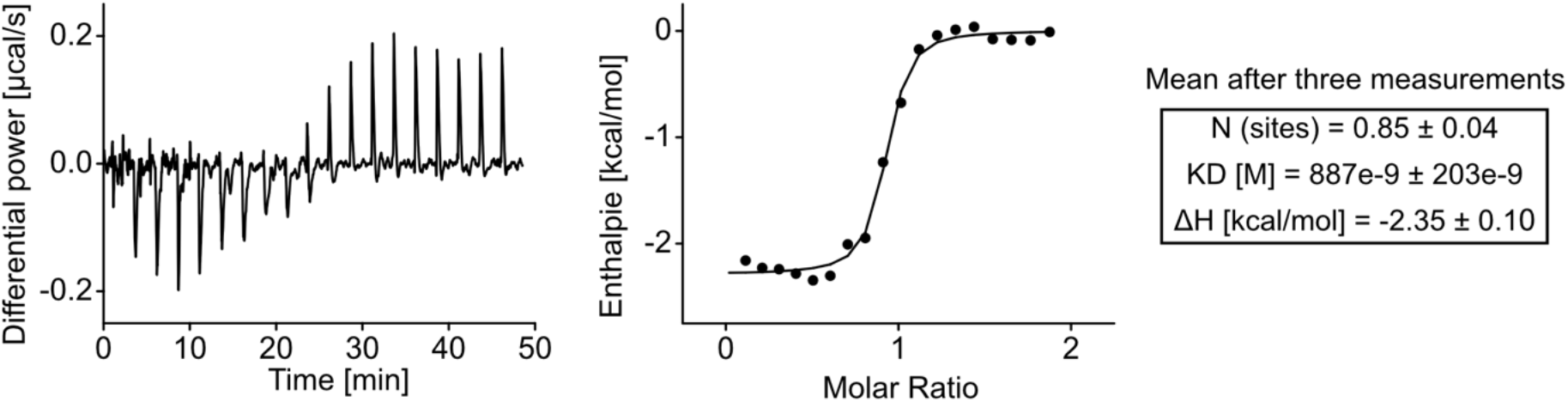
Characterization of HiSiaP/VHH_P_1 binding via ITC. Isothermal titration calorimetry experiment between HiSiaP and the HiSiaP-VHH (left) and binding curve (middle). The thermodynamic parameters from the experiment are shown on the right.

**Fig. S10.**
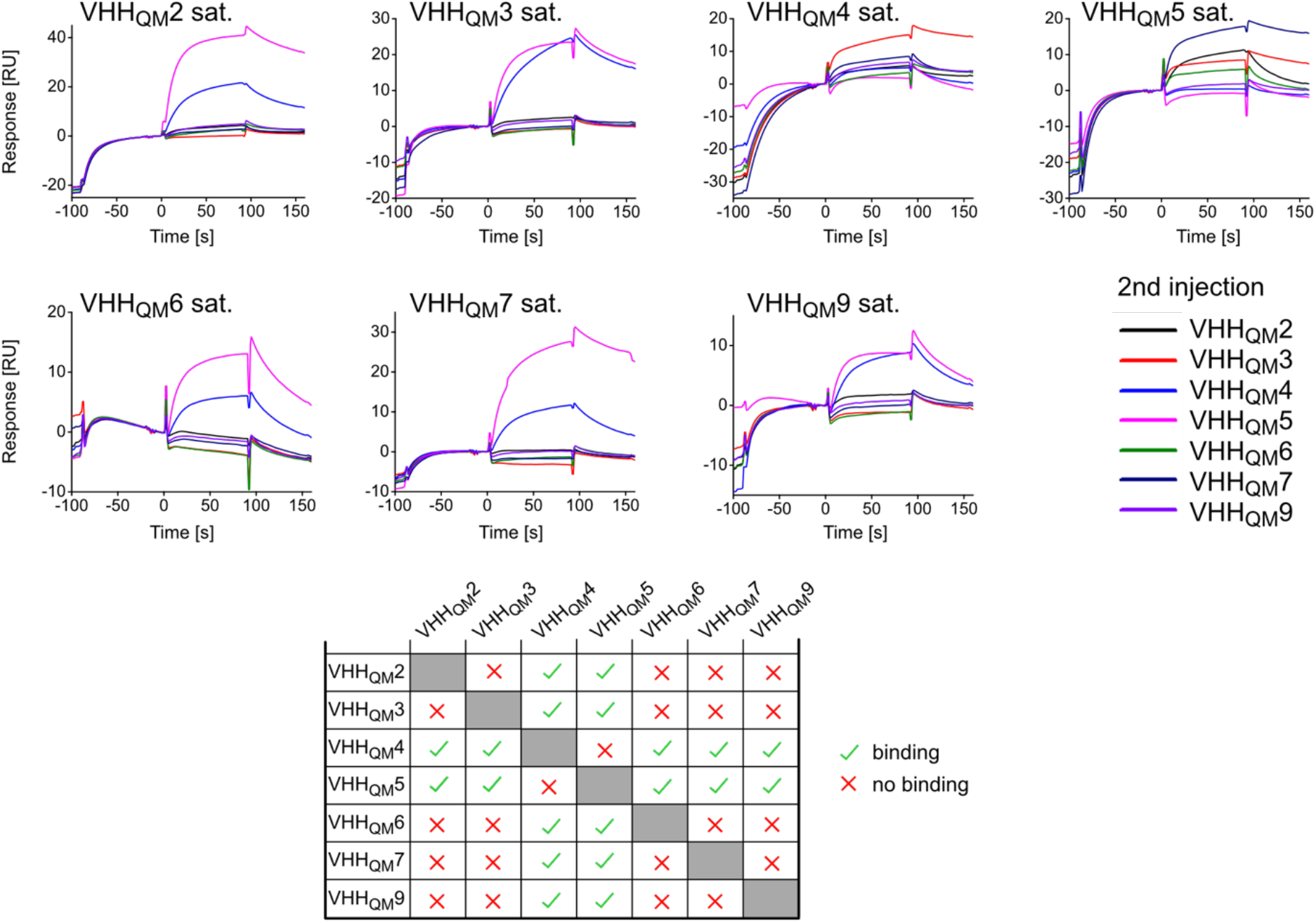
SPR epitope binning. HiSiaQM/VHH interaction was analysed in a competitive binding experiment in which the TRAP transporter (HiSiaQM K273C-biotin), immobilized on an SPR chip, was saturated with a first VHH and the binding behavior of a second VHH was observed. The primary VHH is indicated next to the chromatogram and the secondary VHH is given by the color code. Bottom: Matrix summarizing the epitope binding results.

**Fig. S11.**
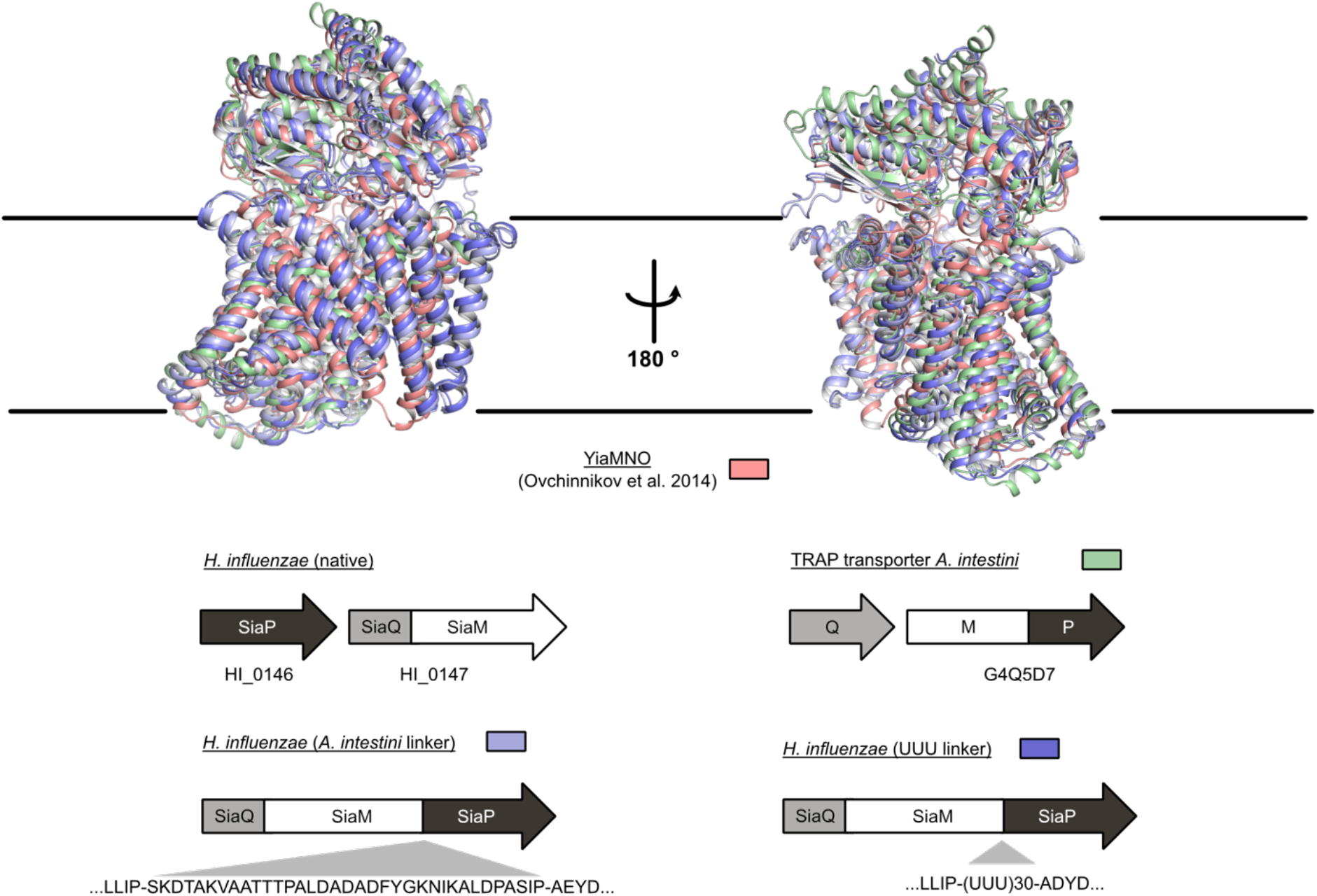
Comparison of models for the tripartite complex. Four different models of the tripartite complex aligned onto each other. The colors are indicated below, next to the organization of the corresponding genes. If domains were fused for modelling, the linkers are specified next to genes. Confidence scores for the different AlphaFold models are shown in Figure S3.

**Fig. S12.**
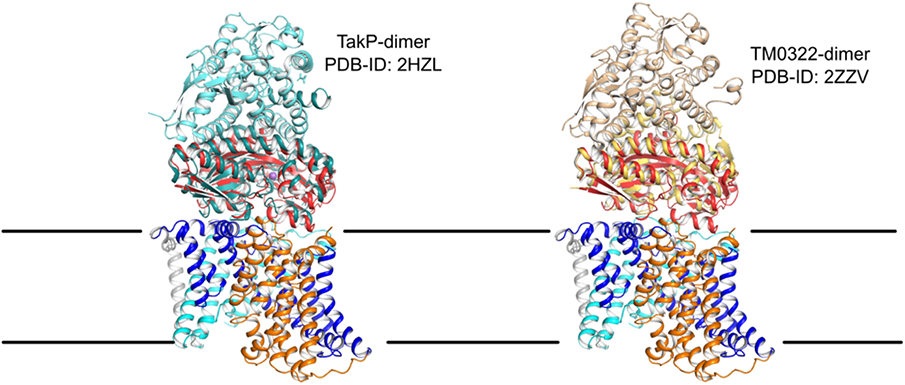
Orientation of dimeric SBPs in the model of the tripartite complex. One chain of the two different dimeric SBPs, TakP (green-blue) and TM0322 (yellow-wheat), was aligned on the HiSiaP SBP (red) in the tripartite complex as predicted by AlphaFold (Figure 3).

**Fig. S13.**
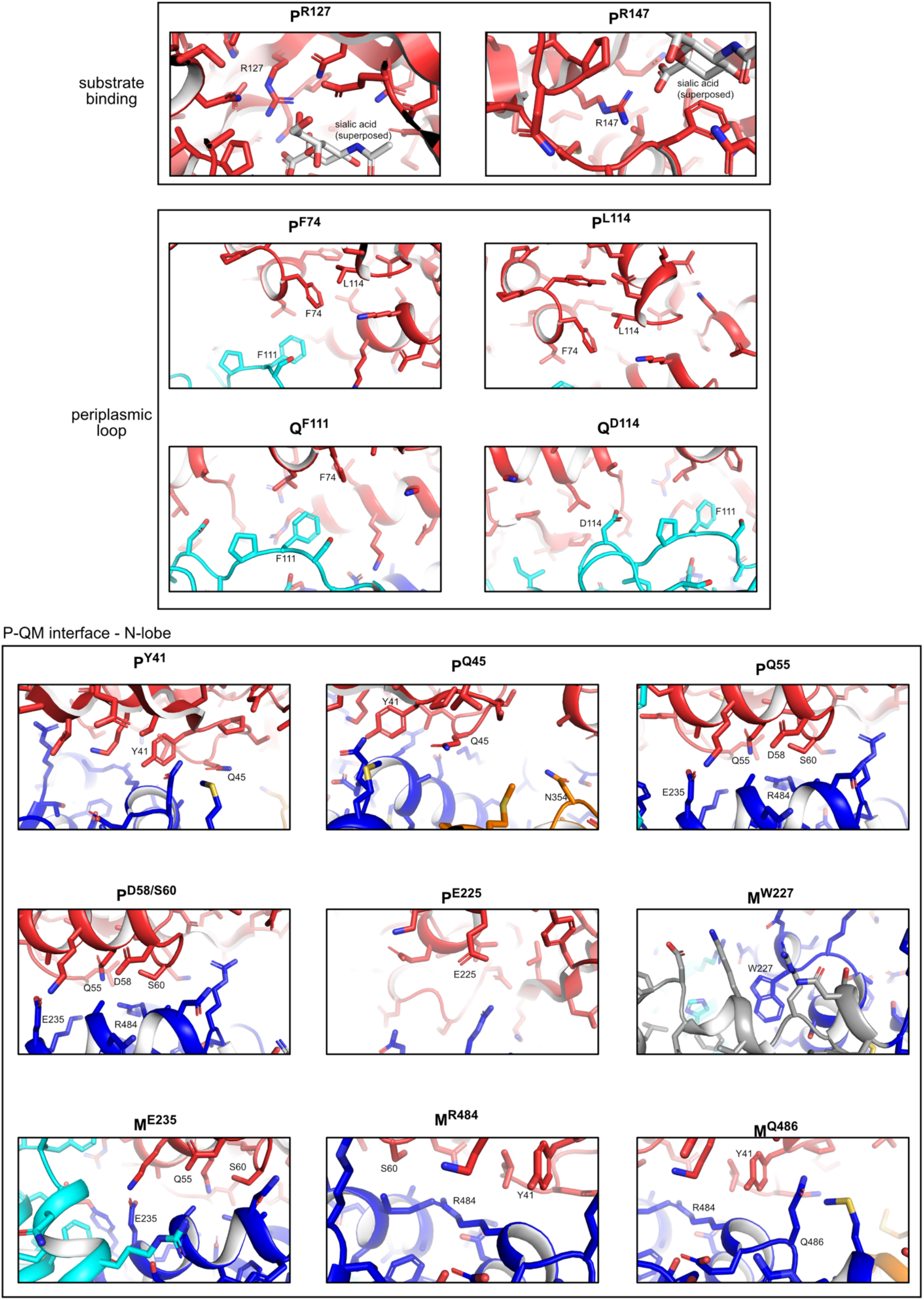
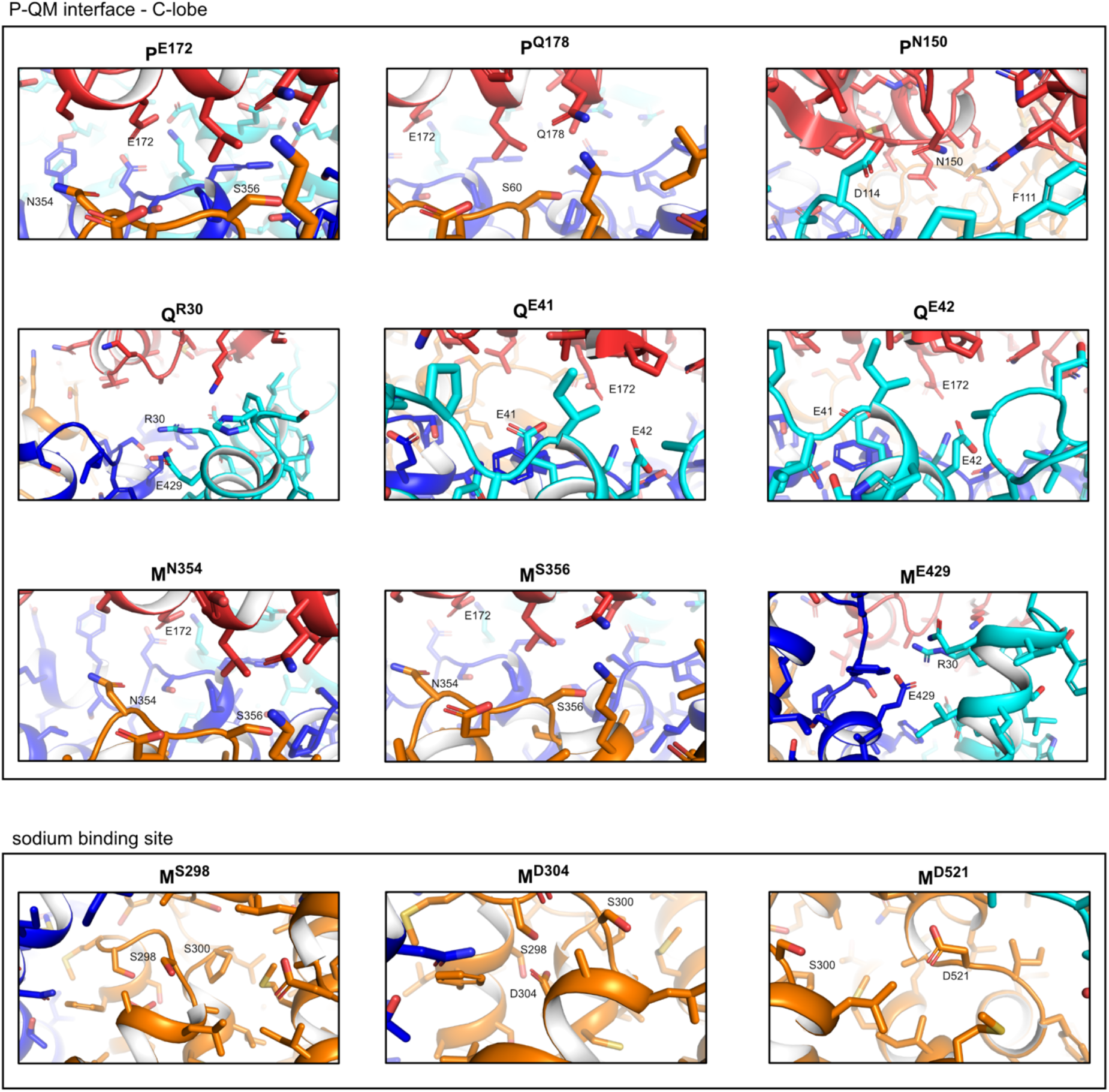
Close-up views of mutants from Figure 3. All residues that were analyzed in the complementation assay (Figure 3) in the main text are shown in detail in the AlphaFold model of the tripartite complex.

**Fig. S14.**
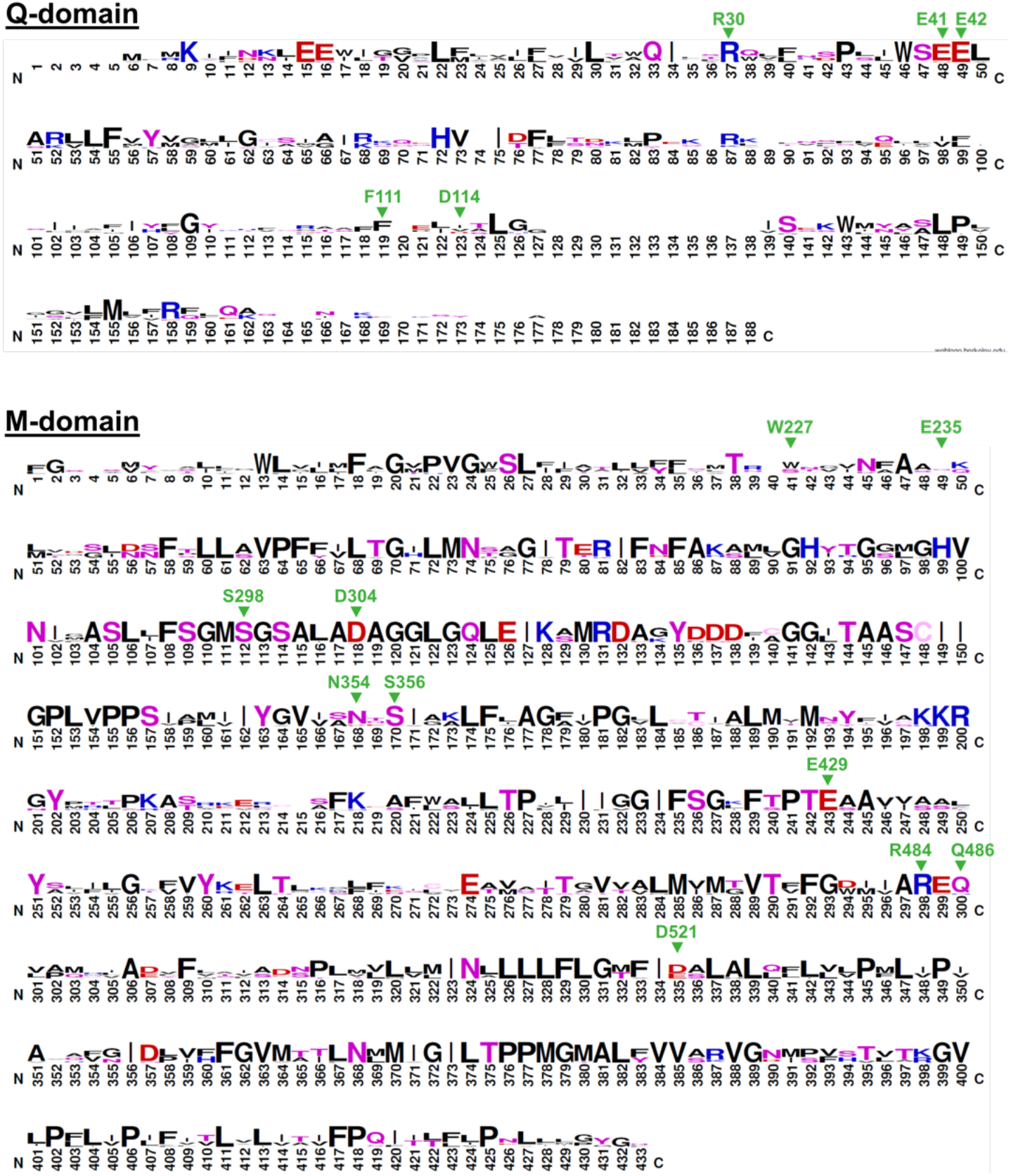
Sequence alignment of sialic acid TRAP transporters. Positions in HiSiaQM that were mutated and analysed in the complementation assay in Figure 3 are specified in green with the HiSiaQM numbering indicated. Sequences (see below) were taken from Chowdhury et al. ^3^ and Vetting et al. ^4^, with just using those with known ligands. Here, just sialic acid bound TRAP transporters are shown, an alignment of TRAP transporters which do not bind sialic acid is shown in Figure S15. The QM sequences were split into Q and M for the alignment using HiSiaQM as a guide. The sequences were aligned using ClustalO^7^ in Jalview^8^ with default settings and sequence logo motifs generated using WebLogo ^5^. SiaQ: NMPREF0198_0032-SiaQ/1-152; NT05HA_0543-SiaQ/1-153; FN1473-SiaQ/1-153; GGC_1730-SiaQ/1-153; SiaQ-HI/1-153; UMN179_02202-SiaQ/1-156; Trebr_2165-SiaQ/1-157; PM1708-SiaQ/1-160; VV2_0733-SiaQ/1-168; PCNPT3_02946-SiaQ/1-169; P3TCK_01175-SiaQ/1-170; VPMS16_3066-SiaQ/1-173; VC1779-SiaQ/1-173; SiaM: Trebr_2166-SiaM/1-427; P3TCK_01175-SiaM/1-427; VV2_0733-SiaM/1-427; VPMS16_3066-SiaM/1-427; PCNPT3_02946-SiaM/1-427; VC1779-SiaM/1-427; NMPREF0198_0032-SiaM/1-429; PM1708-SiaM/1-429; SiaM-HI/1-429; GGC_1730-SiaM/1-429; NT05HA_0543-SiaM/1-429; FN1473-SiaM/1-430; UMN179_02202-SiaM/1-430

**Fig. S15.**
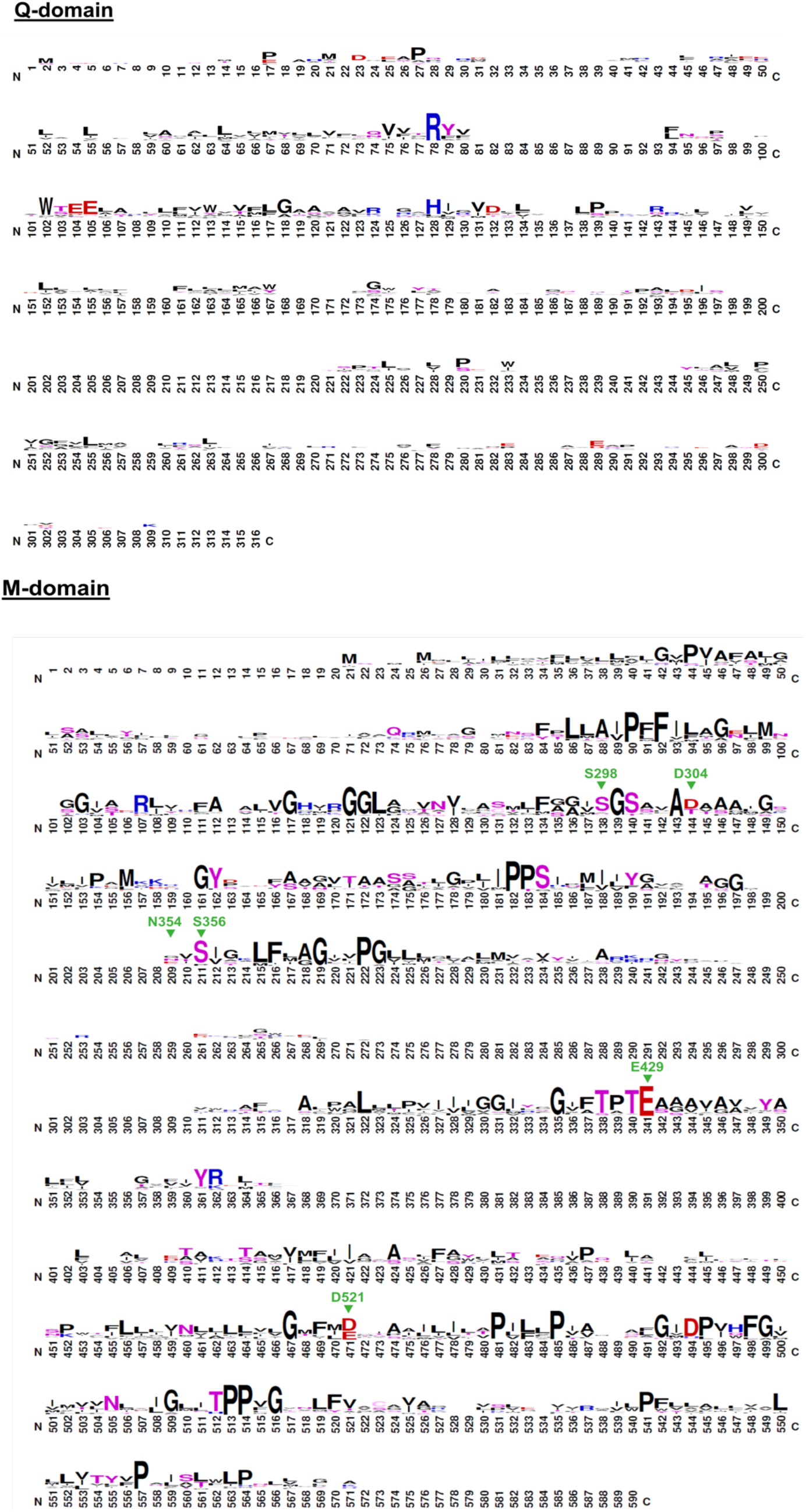
Sequence alignment of TRAP transporters. Positions in HiSiaQM that were mutated and analysed in the complementation assay in Figure 3 are specified in green with the HiSiaQM numbering. As for Figure S14, sequences (see below) were taken from Chowdhury et al. ^3^ and Vetting et al. ^4^, with just using those with known ligands. In contrast to Figure S14, this alignment shows TRAP transporters which are not bind sialic acid. The QM sequences were split into Q and M for alignment using HiSiaQM as a guide. The sequences were aligned using ClustalO ^7^ in Jalview ^8^ with default settings and sequence logo motifs generated using WebLogo ^6^. Q-domains: FN1258-SiaQ/1-147; EFER_1530-SiaQ/1-158; Clobol_03198-SiaQ/1-158; Bpro_3107-SiaQ/1-162; BB2442-SiaQ/1-163; Asuc_0158-SiaQ/1-163; Bpro_1871-SiaQ/1-164; NAS141_03721-SiaQ/1-164; RD1_1052-SiaQ/1-164; HICG_00826-SiaQ/1-165; Rfer_1840-SiaQ/1-165; Dde_0634-SiaQ/1-165; Desal_0342-SiaQ/1-165; BB3421-SiaQ/1-166; NAS141_03681-SiaQ/1-169; Csal_0660-SiaQ/1-169; pro_4736-SiaQ/1-172; SM_b20295-SiaQ/1-172; Desal_2161-SiaQ/1-172; Smed_3836-SiaQ/1-172; CPS_0129-SiaQ/1-173; Oant_3902-SiaQ/1-173; BB0719-SiaQ/1-174; Dde_1548-SiaQ/1-174; RD1_3994-SiaQ/1-175; Csal_2479-SiaQ/1-177; Veis_3954-SiaQ/1-178; RD1_0742-SiaQ/1-180; Oant_4429-SiaQ/1-180; BBta_0128-SiaQ/1-181; Apre_1383-SiaQ/1-183; MIM_c39430-SiaQ/1-185; RPB_3329-SiaQ/1-186; Xaut_3368-SiaQ/1-187; Sdel_0447-SiaQ/1-187; Bpro_0088-SiaQ/1-188; SM_b20442-SiaQ/1-190; Atu3253-SiaQ/1-190; bll6834-SiaQ/1-192; SPO1773-SiaQ/1-192; Bamb_6123-SiaQ/1-192; Csal_0678-SiaQ/1-198; H16_A1328-SiaQ/1-200; RPB_2686-SiaQ/1-201; Shew_1446-SiaQ/1-212; SO_3134-SiaQ/1-219 M-domains: SPO1773-SiaM/1-419;>HICG_00826-SiaM/1-423; Oant_3902-SiaM/1-424; Bpro_1871-SiaM/1-425; Veis_3954-SiaM/1-425; BB3421-SiaM/1-425; Desal_2161-SiaM/1-425; Oant_4429-SiaM/1-425; RD1_1052-SiaM/1-425; Xaut_3368-SiaM/1-426; CPS_0129-SiaM/1-426; RPB_2686-SiaM/1-426; MIM_c39430-SiaM/1-426; Desal_0342-SiaM/1-426; NAS141_03681-SiaM/1-426; Smed_3836-SiaM/1-426; SM_b20295-SiaM/1-426; Csal_0660-SiaM/1-426; Csal_2479-SiaM/1-426; RD1_3994-SiaM/1-427; Dde_1548-SiaM/1-427; Sdel_0447-SiaM/1-427; FN1258-SiaM/1-428; Bpro_3107-SiaM/1-429; Asuc_0158-SiaM/1-430; BB0719-SiaM/1-430; BB2442-SiaM/1-430; BBta_0128-SiaM/1-430; RPB_3329-SiaM/1-430; RD1_0742-SiaM/1-431; Dde_0634-SiaM/1-433; Apre_1383-SiaM/1-433; H16-A1328-SiaM/1-434; Clobol_03198-SiaM/1-435; NAS141_03721-SiaM/1-436; Csal_0678-SiaM/1-440; Bpro_4736-SiaM/1-440; Bpro_0088-SiaM/1-446; Atu3253-SiaM/1-447; Rfer_1840-SiaM/1-449; Bamb_6123-SiaM/1-450; EFER_1530-SiaM/1-452; Shew_1446-SiaM/1-465; SO_3134-SiaM/1-465; bll6834-SiaM/1-468; SM_b20442-SiaM/1-468;

**Fig. S16.**
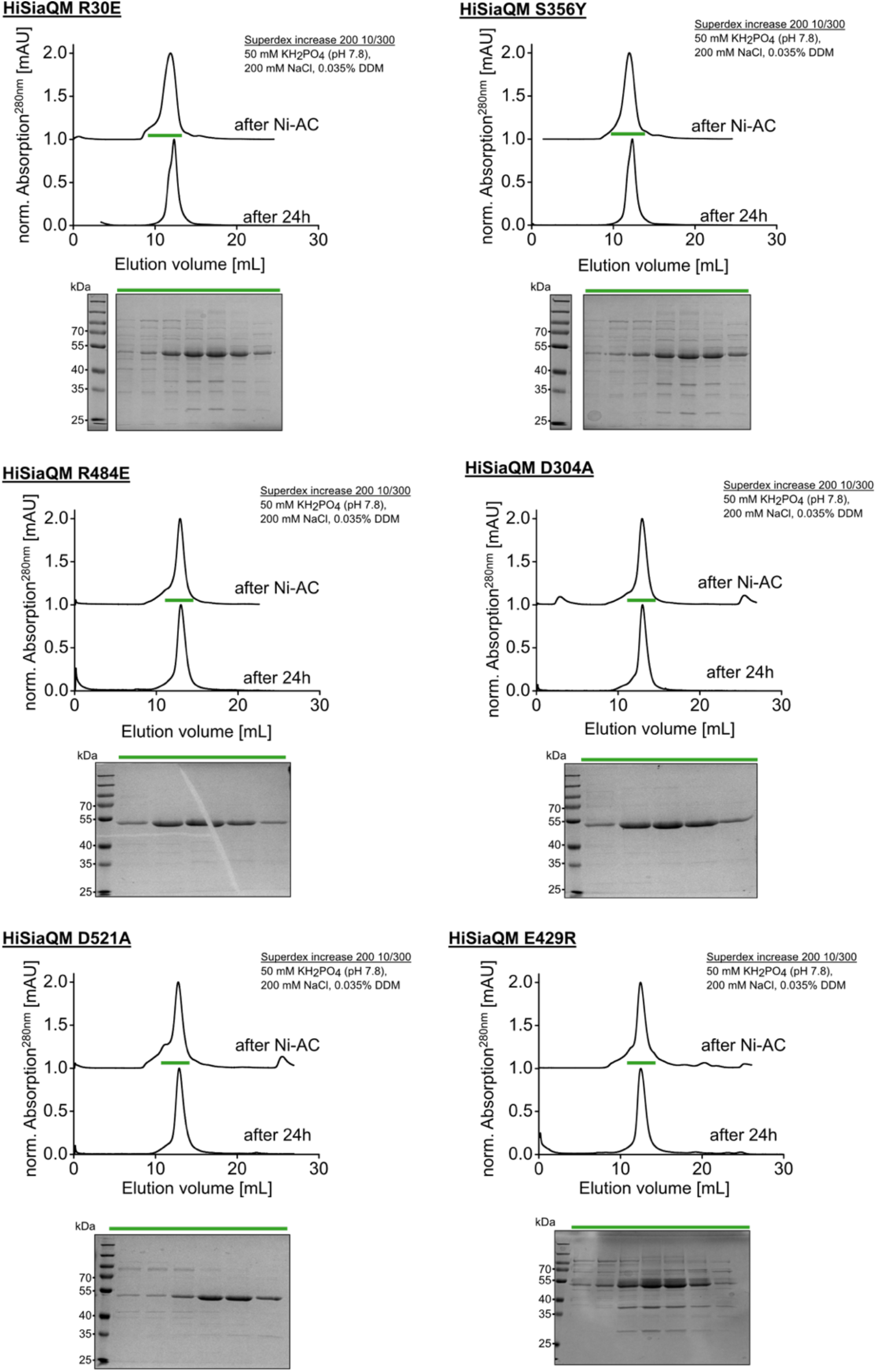
Purification of HiSiaQM mutants. SEC elution profiles of HiSiaQM mutants causing complete loss of function in the complementation assay. The SEC runs were performed using the standard overexpression and purification protocol as described in the methods. Eluted fractions were stored for 24 hours at 4 °C and a further SEC run was performed to assess aggregation. Fractions from the first SEC runs were analyzed by SDS-PAGE, shown next to the chromatograms.

**Fig. S17.**
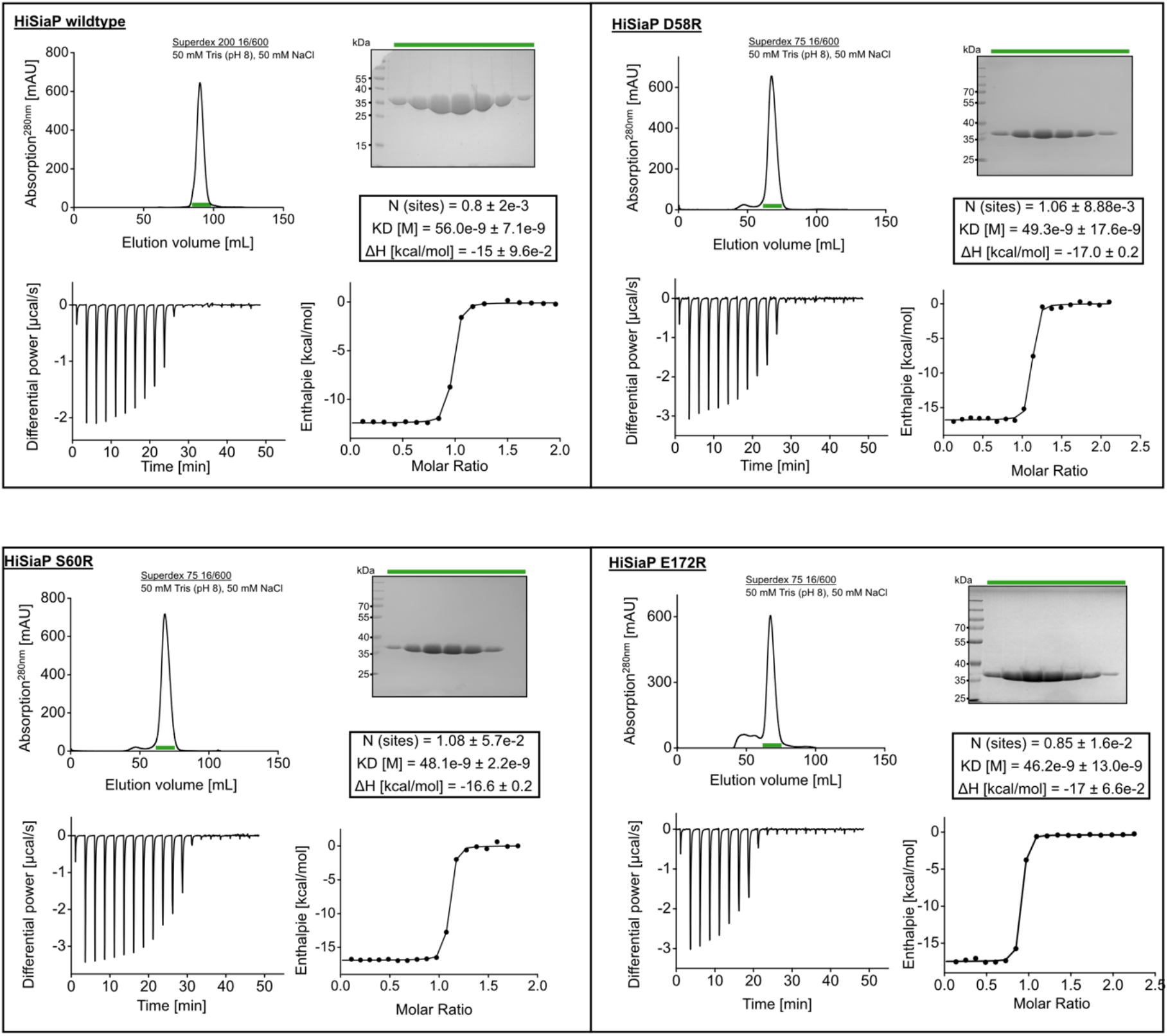
Sialic acid binding of HiSiaP wildtype and mutants. SEC elution profiles of the HiSiaP wildtype and mutants that showed a growth defect in the complementation assay. The SEC runs were performed using the standard overexpression and purification protocol as described in the methods. Eluted fractions were checked via SDS-PAGE, shown below the corresponding chromatograms. The binding of the HiSiaP mutants to sialic acid was analyzed with isothermal titration calorimetry experiments. An exemplary differential power graph and a binding curve is shown for each mutant. The mean binding parameters from three measurements for each mutant are specified.

**Table S1.**
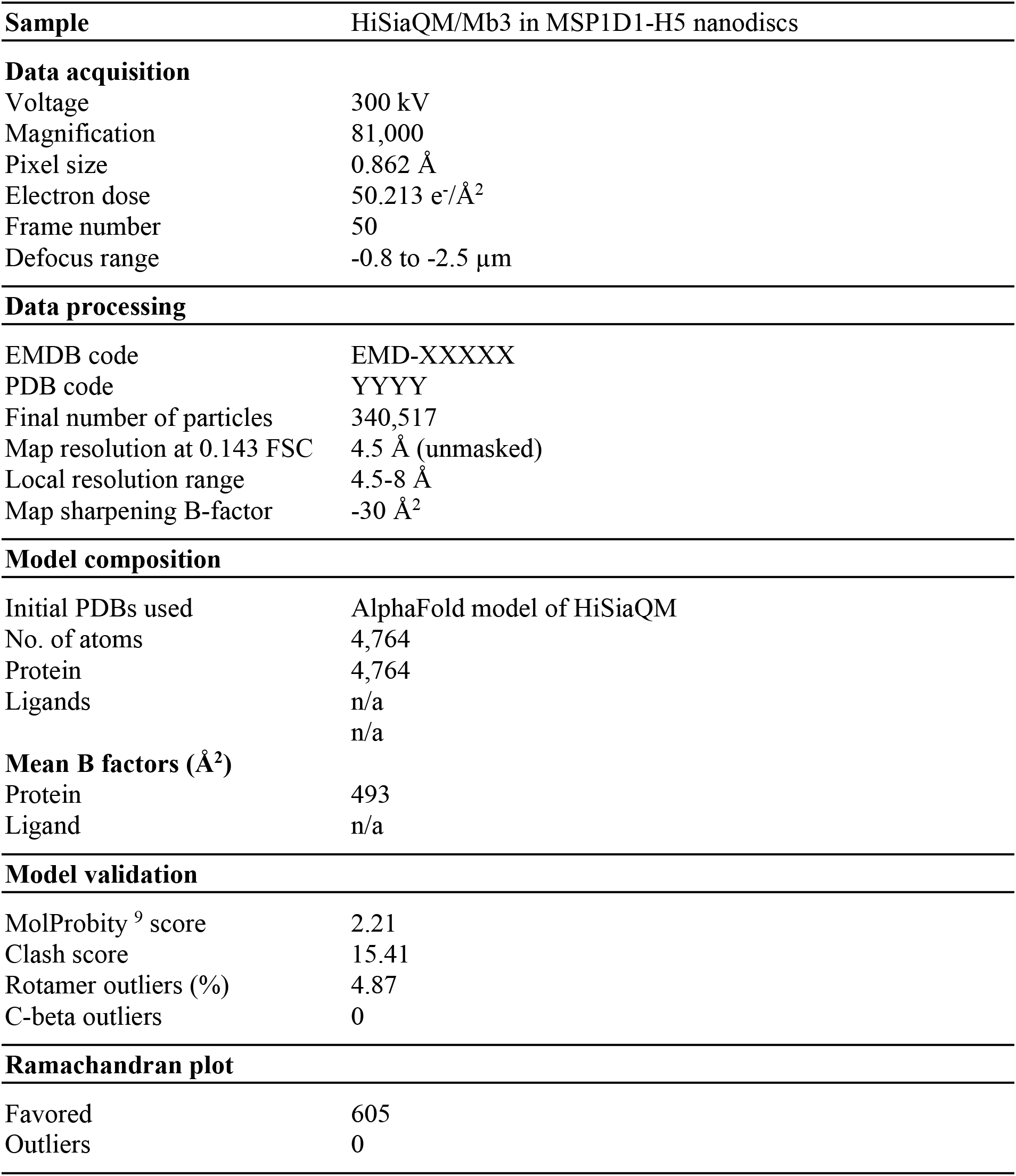
Cryo-EM data collection and processing.

**Table S2.**
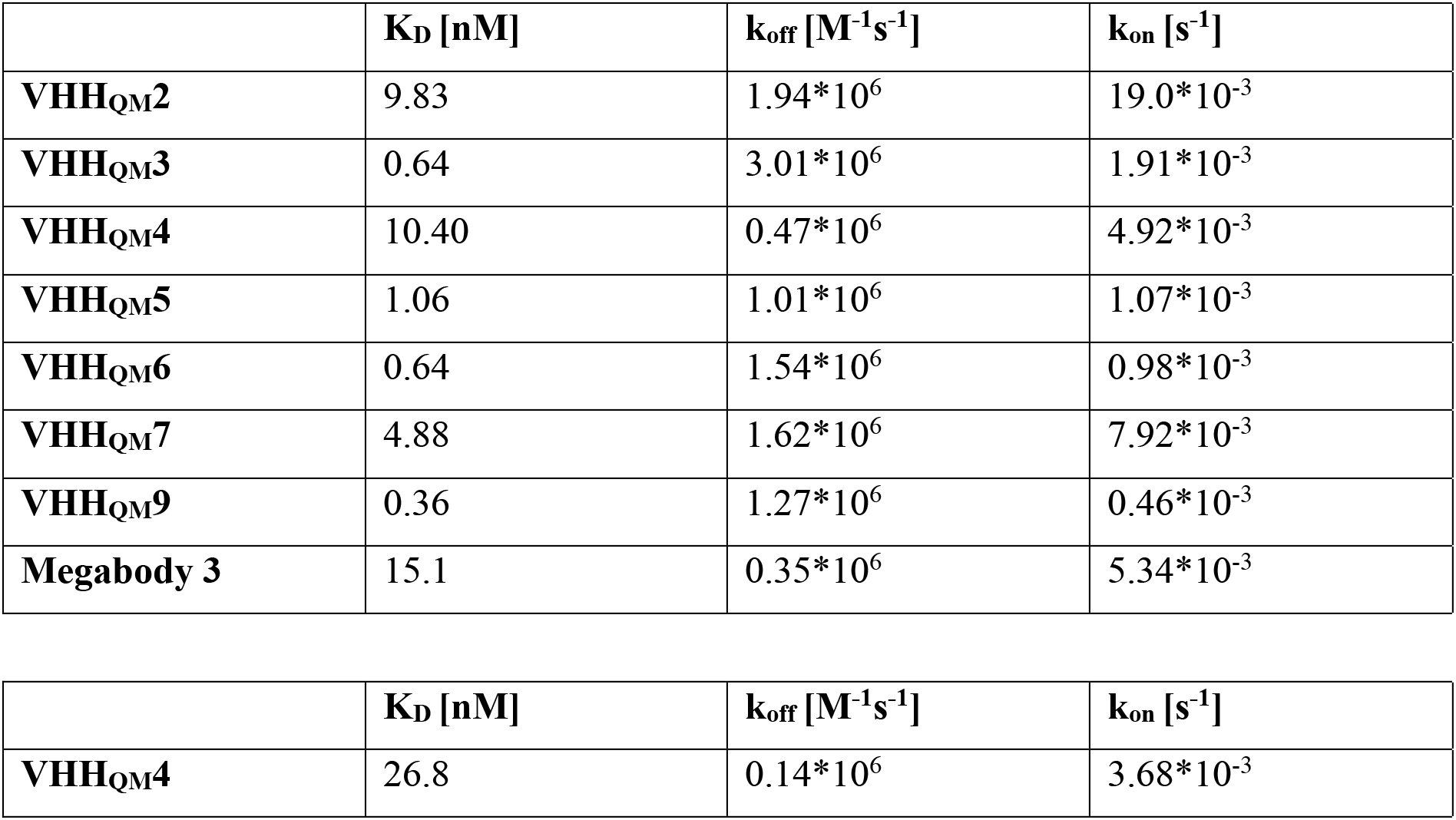
SPR binding parameters of VHHs and megabody on K273C-biotin (top) and E235C-biotin (bottom).

## References

1. Forward, J. A., Behrendt, M. C., Wyborn, N. R., Cross, R. & Kelly, D. J. TRAP transporters: a new family of periplasmic solute transport systems encoded by the dctPQM genes of Rhodobacter capsulatus and by homologs in diverse gram-negative bacteria. Journal of Bacteriology 179, 5482–5493 (1997).

2. Rabus, R., Jack, D. L., Kelly, D. J. & Milton H Saier, J. TRAP transporters: an ancient family of extracytoplasmic solute-receptor-dependent secondary active transporters. Microbiology (Reading, England) 145, 3431–3445 (1999).

3. Rosa, L. T., Bianconi, M. E., Thomas, G. H. & Kelly, D. J. Tripartite ATP-Independent Periplasmic (TRAP) Transporters and Tripartite Tricarboxylate Transporters (TTT): From Uptake to Pathogenicity. Frontiers in cellular and infection microbiology 8, 33 (2018).

4. Mulligan, C., Leech, A. P., Kelly, D. J. & Thomas, G. H. The membrane proteins SiaQ and SiaM form an essential stoichiometric complex in the sialic acid tripartite ATP-independent periplasmic (TRAP) transporter SiaPQM (VC1777-1779) from Vibrio cholerae. J Biol Chem 287, 3598–3608 (2012).

5. Severi, E., Hood, D. W. & Thomas, G. H. Sialic acid utilization by bacterial pathogens. Microbiology 153, 2817–2822 (2007).

6. Erkens, G. B., Hänelt, I., Goudsmits, J. M. H., Slotboom, D. J. & van Oijen, A. M. Unsynchronised subunit motion in single trimeric sodium-coupled aspartate transporters. Nature 502, 119–123 (2013).

7. Mancusso, R., Gregorio, G. G., Liu, Q. & Wang, D.-N. Structure and mechanism of a bacterial sodium-dependent dicarboxylate transporter. Nature 491, 622–626 (2012).

8. Jumper, J. et al. Highly accurate protein structure prediction with AlphaFold. Nature 596, 583–589 (2021).

9. Mulligan, C., Kelly, D. J. & Thomas, G. H. Tripartite ATP-independent periplasmic transporters: application of a relational database for genome-wide analysis of transporter gene frequency and organization. J Mol Microbiol Biotechnol 12, 218–226 (2007).

10. Kelly, D. J. & Thomas, G. H. The tripartite ATP-independent periplasmic (TRAP) transporters of bacteria and archaea. FEMS Microbiology Reviews 25, 405–424 (2001).

11. Mulligan, C., Fischer, M. & Thomas, G. H. Tripartite ATP-independent periplasmic (TRAP) transporters in bacteria and archaea. FEMS Microbiology Reviews 35, 68–86 (2011).

12. Winnen, B., Hvorup, R. N. & Saier, M. H. The tripartite tricarboxylate transporter (TTT) family. Research in Microbiology 154, 457–465 (2003).

13. Mulligan, C. et al. The substrate-binding protein imposes directionality on an electrochemical sodium gradient-driven TRAP transporter. Proceedings of the National Academy of Sciences of the United States of America 106, 1778–1783 (2009).

14. Severi, E. et al. Sialic acid transport in Haemophilus influenzae is essential for lipopolysaccharide sialylation and serum resistance and is dependent on a novel tripartite ATP-independent periplasmic transporter. Molecular Microbiology 58, 1173–1185 (2005).

15. Chowdhury, N., et al. The VC1777–VC1779 proteins are members of a sialic acid-specific subfamily of TRAP transporters (SiaPQM) and constitute the sole route of sialic acid uptake in the human pathogen Vibrio cholerae. Microbiology 158, 2158–2167 (2012).

16. Allen, S., Zaleski, A., Johnston, J. W., Gibson, B. W. & Apicella, M. A. Novel sialic acid transporter of Haemophilus influenzae. Infection and Immunity 73, 5291–5300 (2005).

17. Bouchet, V. et al. Host-derived sialic acid is incorporated into Haemophilus influenzae lipopolysaccharide and is a major virulence factor in experimental otitis media. Proceedings of the National Academy of Sciences of the United States of America 100, 8898–8903 (2003).

18. Jenkins, G. A. et al. Sialic acid mediated transcriptional modulation of a highly conserved sialometabolism gene cluster in Haemophilus influenzae and its effect on virulence. BMC Microbiol 10, 48 (2010).

19. Almagro-Moreno, S. & Boyd, E. F. Sialic acid catabolism confers a competitive advantage to pathogenic vibrio cholerae in the mouse intestine. Infection and immunity 77, 3807–3816 (2009).

20. Beyer, P. & Paulin, S. Priority pathogens and the antibiotic pipeline: an update. Bulletin of the World Health Organization 98, 151 (2020).

21. Locher, K. P. Structure and mechanism of ATP-binding cassette transporters. Philosophical Transactions of the Royal Society B: Biological Sciences 364, 239–245 (2009).

22. Müller, A. et al. Conservation of structure and mechanism in primary and secondary transporters exemplified by SiaP, a sialic acid binding virulence factor from Haemophilus influenzae. Journal of Biological Chemistry 281, 22212–22222 (2006).

23. Vetting, M. W. et al. Experimental strategies for functional annotation and metabolism discovery: targeted screening of solute binding proteins and unbiased panning of metabolomes. Biochemistry 54, 909–931 (2015).

24. Gangi Setty, T., Cho, C., Govindappa, S., Apicella, M. A. & Ramaswamy, S. Bacterial periplasmic sialic acid-binding proteins exhibit a conserved binding site. Acta Crystallographica Section D 70, 1801–1811 (2014).

25. Peter, M. F. et al. Triggering closure of a sialic acid TRAP transporter substrate binding protein through binding of natural or artificial substrates. Journal of Molecular Biology 433, 166756 (2021).

26. Johnston, J. W. et al. Characterization of the N-acetyl-5-neuraminic acid-binding site of the extracytoplasmic solute receptor (SiaP) of nontypeable Haemophilus influenzae strain 2019. Journal of Biological Chemistry 283, 855–865 (2008).

27. Herrou, J. et al. Structure-based mechanism of ligand binding for periplasmic solute-binding protein of the Bug family. Journal of Molecular Biology 373, 954–964 (2007).

28. Berntsson, R. P. A., Smits, S. H. J., Schmitt, L., Slotboom, D.-J. & Poolman, B. A structural classification of substrate-binding proteins. FEBS Letters 584, 2606–2617 (2010).

29. Fischer, M. et al. Tripartite ATP-independent Periplasmic (TRAP) Transporters Use an Arginine-mediated Selectivity Filter for High Affinity Substrate Binding. J Biol Chem 290, 27113–27123 (2015).

30. Darby, J. F. et al. Water networks can determine the affinity of ligand binding to proteins. Journal of the American Chemical Society 141, 15818–15826 (2019).

31. Glaenzer, J., Peter, M. F., Thomas, G. H. & Hagelueken, G. PELDOR Spectroscopy Reveals Two Defined States of a Sialic Acid TRAP Transporter SBP in Solution. Biophysical Journal 112, 109–120 (2017).

32. Uchański, T. et al. Megabodies expand the nanobody toolkit for protein structure determination by single-particle cryo-EM. Nat Methods 18, 60–68 (2021).

33. Mulligan, C. et al. The bacterial dicarboxylate transporter VcINDY uses a two-domain elevator-type mechanism. Nat Struct Mol Biol 23, 256–263 (2016).

34. Vergara-Jaque, A., Fenollar-Ferrer, C., Mulligan, C., Mindell, J. A. & Forrest, L. R. Family resemblances: A common fold for some dimeric ion-coupled secondary transporters. J Gen Physiol 146, 423–434 (2015).

35. Denisov, I. G., Grinkova, Y. V., Lazarides, A. A. & Sligar, S. G. Directed self-assembly of monodisperse phospholipid bilayer Nanodiscs with controlled size. Journal of the American Chemical Society 126, 3477–3487 (2004).

36. Puthenveetil, R., Nguyen, K. & Vinogradova, O. Nanodiscs and solution NMR: preparation, application and challenges. Nanotechnology reviews 6, 111–125 (2017).

37. Punjani, A., Rubinstein, J. L., Fleet, D. J. & Brubaker, M. A. cryoSPARC: algorithms for rapid unsupervised cryo-EM structure determination. Nat Methods 14, 290–296 (2017).

38. Cramer, P. AlphaFold2 and the future of structural biology. Nat Struct Mol Biol 28, 704–705 (2021).

39. Liebschner, D. et al. Macromolecular structure determination using X-rays, neutrons and electrons: recent developments in Phenix. Acta Crystallographica Section D: Structural Biology 75, 861–877 (2019).

40. Ovchinnikov, S., Kamisetty, H. & Baker, D. Robust and accurate prediction of residue– residue interactions across protein interfaces using evolutionary information. eLife 3, 1061–1021 (2014).

41. Vergara-Jaque, A., Fenollar-Ferrer, C., Kaufmann, D. & Forrest, L. R. Repeat-swap homology modeling of secondary active transporters: updated protocol and prediction of elevator-type mechanisms. Front Pharmacol 6, 183 (2015).

42. Crisman, T. J., Qu, S., Kanner, B. I. & Forrest, L. R. Inward-facing conformation of glutamate transporters as revealed by their inverted-topology structural repeats. Proceedings of the National Academy of Sciences of the United States of America 106, 20752–20757 (2009).

43. Radestock, S. & Forrest, L. R. The alternating-access mechanism of MFS transporters arises from inverted-topology repeats. Journal of Molecular Biology 407, 698–715 (2011).

44. Mulligan, C. & Mindell, J. A. Pinning Down the Mechanism of Transport: Probing the Structure and Function of Transporters Using Cysteine Cross-Linking and Site-Specific Labeling. Methods in Enzymology 594, 165–202 (2017).

45. Emsley, P. & Cowtan, K. Coot: model-building tools for molecular graphics. Acta Crystallographica Section D 60, 2126–2132 (2004).

46. Trinco, G. et al. Kinetic mechanism of Na+-coupled aspartate transport catalyzed by GltTk. Commun Biol 4, 751 (2021).

47. Schenck, S. et al. Generation and Characterization of Anti-VGLUT Nanobodies Acting as Inhibitors of Transport. Biochemistry 56, 3962–3971 (2017).

48. Mireku, S. A., Sauer, M. M., Glockshuber, R. & Locher, K. P. Structural basis of nanobody-mediated blocking of BtuF, the cognate substrate-binding protein of the Escherichia coli vitamin B12 transporter BtuCD. Sci Rep 7, 14296 (2017).

49. Errasti-Murugarren, E. et al. L amino acid transporter structure and molecular bases for the asymmetry of substrate interaction. Nature communications 10, 1–12 (2019).

50. Severi, E., Hosie, A. H. F., Hawkhead, J. A. & Thomas, G. H. Characterization of a novel sialic acid transporter of the sodium solute symporter (SSS) family and in vivo comparison with known bacterial sialic acid transporters. FEMS Microbiology Letters 304, 47–54 (2010).

51. Cash, P., Argo, E. & Bruce, K. D. Characterisation of Haemophilus influenzae proteins by two-dimensional gel electrophoresis. Electrophoresis 16, 135–148 (1995).

52. Kolker, E. et al. Initial proteome analysis of model microorganism Haemophilus influenzae strain Rd KW20. Journal of Bacteriology 185, 4593–4602 (2003).

53. Krissinel, E. & Henrick, K. Inference of macromolecular assemblies from crystalline state. Journal of Molecular Biology 372, 774–797 (2007).

54. Gonin, S. et al. Crystal structures of an Extracytoplasmic Solute Receptor from a TRAP transporter in its open and closed forms reveal a helix-swapped dimer requiring a cation for alpha-keto acid binding. BMC Struct Biol 7, 11 (2007).

55. Akiyama, N., Takeda, K. & Miki, K. Crystal structure of a periplasmic substrate-binding protein in complex with calcium lactate. Journal of Molecular Biology 392, 559–565 (2009).

56. Oldham, M. L., Khare, D., Quiocho, F. A., Davidson, A. L. & Chen, J. Crystal structure of a catalytic intermediate of the maltose transporter. Nature 450, 515–521 (2007).

57. Wahlgren, W. Y. et al. Substrate-bound outward-open structure of a Na^+^-coupled sialic acid symporter reveals a new Na^+^ site. Nat Commun 9, 1753 (2018).

58. Marinelli, F. et al. Evidence for an allosteric mechanism of substrate release from membrane-transporter accessory binding proteins. Proceedings of the National Academy of Sciences of the United States of America 108, E1285–E1292 (2011).

59. Pettersen, E. F. et al. UCSF ChimeraX: Structure visualization for researchers, educators, and developers. Protein Sci 30, 70–82 (2021).

60. Waterhouse, A. M., Procter, J. B., Martin, D. M. A., Clamp, M. & Barton, G. J. Jalview Version 2—a multiple sequence alignment editor and analysis workbench. Bioinformatics 25, 1189–1191 (2009).

61. Celniker, G. et al. ConSurf: Using Evolutionary Data to Raise Testable Hypotheses about Protein Function. Israel Journal of Chemistry 53, 199–206 (2013).

62. Liu, H. & Naismith, J. H. An efficient one-step site-directed deletion, insertion, single and multiple-site plasmid mutagenesis protocol. BMC Biotechnology 8, 91 (2008).

63. Ritchie, T. K. et al. Reconstitution of membrane proteins in phospholipid bilayer nanodiscs. Methods in Enzymology 464, 211–231 (2009).

64. Kalivoda, K. A., Steenbergen, S. M. & Vimr, E. R. Control of the Escherichia coli sialoregulon by transcriptional repressor NanR. Journal of Bacteriology 195, 4689–4701 (2013).

65. Plumbridge, J. & Vimr, E. Convergent pathways for utilization of the amino sugars N-acetylglucosamine, N-acetylmannosamine, and N-acetylneuraminic acid by Escherichia coli. Journal of Bacteriology 181, 47–54 (1999).

66. Plumbridge, J. A. Repression and induction of the nag regulon of Escherichia coli K-12: the roles of nagC and nagA in maintenance of the uninduced state. Mol Microbiol 5, 2053–2062 (1991).

67. Link, A. J., Phillips, D. & Church, G. M. Methods for generating precise deletions and insertions in the genome of wild-type Escherichia coli: application to open reading frame characterization. Journal of Bacteriology 179, 6228–6237 (1997).

## References for supporting information

1. Ovchinnikov, S., Kamisetty, H. & Baker, D. Robust and accurate prediction of residue– residue interactions across protein interfaces using evolutionary information. eLife 3, 1061–1021 (2014).

2. Jumper, J. et al. Highly accurate protein structure prediction with AlphaFold. Nature 596, 583–589 (2021).

3. Chowdhury, N., et al. The VC1777–VC1779 proteins are members of a sialic acid-specific subfamily of TRAP transporters (SiaPQM) and constitute the sole route of sialic acid uptake in the human pathogen Vibrio cholerae. Microbiology 158, 2158–2167 (2012).

4. Vetting, M. W. et al. Experimental strategies for functional annotation and metabolism discovery: targeted screening of solute binding proteins and unbiased panning of metabolomes. Biochemistry 54, 909–931 (2015).

5. Schneider, T. D. & Stephens, R. M. Sequence logos: a new way to display consensus sequences. Nucleic Acids Res 18, 6097–6100 (1990).

6. Crooks, G. E., Hon, G., Chandonia, J.-M. & Brenner, S. E. WebLogo: a sequence logo generator. Genome research 14, 1188–1190 (2004).

7. Sievers, F. & Higgins, D. G. in Multiple sequence alignment methods 105–116 (Springer, 2014).

8. Waterhouse, A. M., Procter, J. B., Martin, D. M. A., Clamp, M. & Barton, G. J. Jalview Version 2—a multiple sequence alignment editor and analysis workbench. Bioinformatics 25, 1189–1191 (2009).

9. Williams, C. J. et al. MolProbity: More and better reference data for improved all-atom structure validation. Protein Science 27, 293–315 (2018).

